# A DNA-guided Argonaute Protein Functions in DNA Replication in *Thermus thermophilus*

**DOI:** 10.1101/869172

**Authors:** Samson M. Jolly, Ildar Gainetdinov, Karina Jouravleva, Han Zhang, Lara Strittmatter, Gregory M. Hendricks, Avantika Dhabaria, Beatrix Ueberheide, Phillip D. Zamore

**Affiliations:** Department of Biochemistry and Molecular Pharmacology, University of Massachusetts Medical School, Worcester, MA 01605, USA; Howard Hughes Medical Institute and RNA Therapeutics Institute, University of Massachusetts Medical School, Worcester, MA 01605, USA; Department of Cell Biology, University of Massachusetts Medical School, Worcester, Massachusetts, USA; Proteomics Laboratory, New York University School of Medicine, New York, NY 10016, USA

## Abstract

Argonaute proteins use nucleic acid guides to protect organisms against transposons and viruses. In the eubacterium *Thermus thermophilus*, the DNA-guided Argonaute TtAgo defends against transformation by DNA plasmids. Here, we report that TtAgo also participates in DNA replication. TtAgo binds small DNA guides derived from the chromosomal region where replication terminates and associates with proteins known to act in DNA replication. *T. thermophilus* deploys a single type II topoisomerase, gyrase. When gyrase is inhibited, *T. thermophilus* relies on TtAgo to complete replication of its circular genome; loss of both gyrase and TtAgo activity produces long filaments that fail to separate into individual bacteria. We propose that the primary role of TtAgo is to help *T. thermophilus* disentangle the catenated circular chromosomes made by DNA replication.

**One Sentence Summary:** The DNA-guided Argonaute protein of *Thermus thermophilus* helps separate daughter chromosomes at the end of DNA replication.

---

In all domains of life, short nucleic acid guides direct Argonaute (AGO) proteins to defend against transposons, viruses, or plasmids. Among sequenced eubacterial genomes, ~17% encode AGO proteins, whose in vivo functions remain poorly understood (*1–3*). Prokaryotic AGOs often reside in genomic neighborhoods populated by genes acting in host defense (*4*). Unlike eukaryotic Argonautes which bind RNA guides, some prokaryotic AGOs use DNA guides to target DNA cleavage. Such DNA-guided, DNA endonucleases include TtAgo (TT_P0026) from *Thermus thermophilus*, a Gram-negative eubacterium that thrives at 65°C, (*5–7*). In vivo, TtAgo reduces susceptibility to transformation by DNA plasmids (*5*), and, when produced in *E. coli*, randomly acquires guide DNAs from plasmids and the genome. *T. thermophilus* encodes its genes on both a large circular chromosome (~1.9 Mb) and one (HB27 strain) or more (HB8 strain) megaplasmids as large as 0.27 Mb (*8–10*). Each cell contains 4–7 copies of the chromosome and megaplasmid, which segregate randomly between daughter cells (*11, 12*). In both HB27 and HB8, TtAgo resides on the megaplasmid with other host defense genes.

In eukaryotes, RNA-guided human and plant AGOs can acquire RNA guides from double-stranded DNA (dsDNA) breaks and act in DNA repair, and human AGO2 associates with Rad51 (*13–15*). In *Drosophila* S2 cells, Ago2 can similarly acquire RNA guides from dsDNA breaks (*16, 17*). That some prokaryotic AGOs bind small DNAs (smDNA) suggests potential roles in replication, repair or recombination. In prokaryotes, smDNA guides may be readily available, at least in *E. coli*, because the repair and recombination complex RecBCD generates smDNAs that integrate as spacers into CRISPR loci via replication-dependent mechanism (*18*). We sought to understand the role of TtAgo in vivo in *T. thermophilus*.

## TtAgo binds smDNA guides from the terminus of replication

In vivo, TtAgo binds 15–18 nt long DNA guides that derive mainly from a 39 kb region on the *T. thermophilus* chromosome, directly opposite the origin of replication (Fig. 1). We used a polyclonal antibody raised against the entire TtAgo protein to immunoprecipitate the nucleic acids associated with TtAgo in wild-type or mutant *T. thermophilus* HB27 strains grown at 65°C (Fig. 1, A to C and fig. S1A and B). Long (>1000 nt) and short (15–18 nt), 5’ monophosphorylated nucleic acids coimmunoprecipitate with wild-type TtAgo (Fig. 1D). Sensitivity to DNase and resistance to RNase identified the nucleic acids associated with TtAgo as DNA. High-throughput sequencing of the long DNAs showed that they mapped essentially to the entire chromosome and megaplasmid (fig. S1C). In contrast, within both the chromosome and megaplasmid, the smDNAs mainly derived from a region 183° clockwise from the annotated origin of replication (*ori*) (Fig. 1E). Among the 15–18 nt sequences mapping just once to the genome (96% of reads), 90% aligned to the chromosome and 10% to the megaplasmid, a distribution comparable to the relative sizes of the two genomic components (88% and 12% of the genome, respectively), indicating that neither is preferentially sampled. As observed for TtAgo expressed in *E. coli* (*5*), the smDNAs coimmunoprecipitated with TtAgo in vivo typically began with cytidine, but otherwise had a GC content identical to the *T. thermophilus* genome (fig. S1, D and E)

**Fig. 1.**
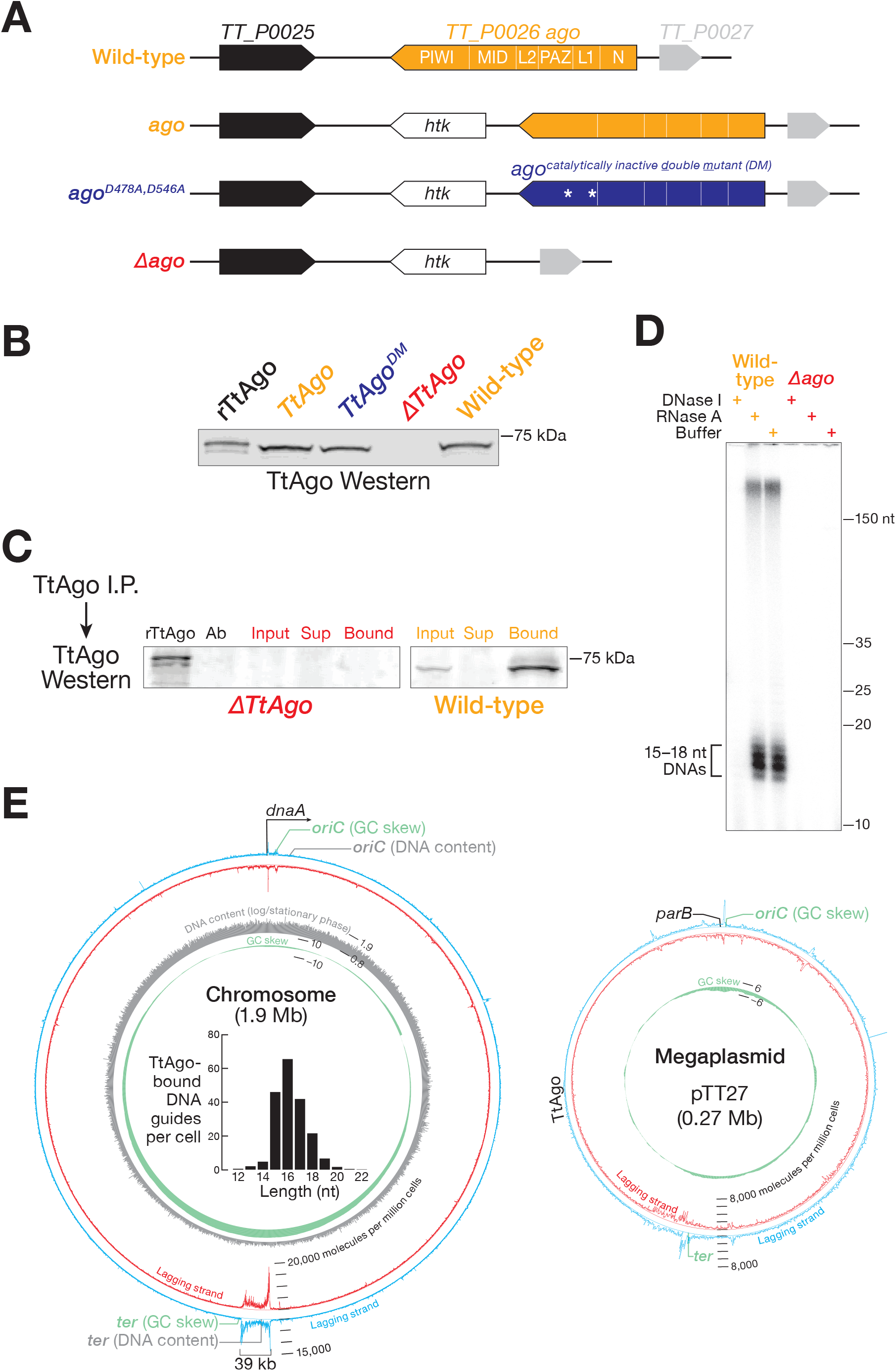
TtAgo expression and TtAgo-bound nucleic acids. (**A**) Study strains: wildtype *T. thermophilus* strain HB27; *ago*, wild-type bearing a thermostable kanamycin resistance gene (*htk*) at the endogenous *ago* locus; *ago^DM^*, D478A, D546A mutant expressing catalytically inactive TtAgo; *Δago*, a deletion mutant lacking the *ago* gene. (**B**) Detection of TtAgo expression at OD_600_ = 0.5. rTtAgo: purified, recombinant TtAgo. (**C**) Immunoprecipitation of TtAgo from lysates of the bacteria in (B). (**D**) DNase and RNase digestion of nucleic acids bound to TtAgo following dephosphorylation and radiolabeling. (**E**) Alignment to the wild-type HB27 genome of TtAgo-bound small DNAs sequenced using a method that requires a 5’ monophosphorylated end. Reads are grouped in 100 bp bins. Inner grey circle: ratio of DNA content between logarithmic and stationary phases. Inner green circle: cumulative GC-skew analysis. Bar graph illustrates length distribution of genome-mapping 5’-phosphorylated DNA guides bound to TtAgo.

That TtAgo binds smDNAs from a ~39 kbp region directly opposite the origin of replication suggests these ssDNA guides arise from a region encompassing the terminus (*ter*), where replication of the circular chromosome and megaplasmid finishes, producing catenated daughter chromosomes that must be separated by topoisomerase II enzymes. Because the terminus has not yet been defined for *T. thermophilus*, we first used GC skew analysis, a method that identifies the origin and terminus of replication based on the distinct nucleotide content of the leading and lagging DNA strands (skew calculated as [G-C]/[G+C]) (*19*). Cumulative GC skew analysis placed the *ori* of the strain HB27 chromosome at 1,541,431 bp, < 2 kbp from the site (1,540,040 bp for our lab strain by long-read sequencing and de novo assembly) deduced from the location of the gene encoding the replication initiating factor *dnaA*, which typically initiates replication by binding a site immediately after its own coding sequence (*20*) (Fig. 1E). Cumulative GC skew analysis located the chromosomal terminus site 187° opposite the *ori* at 626,088 bp, within the region to which the co-immunoprecipitated smDNAs align (Fig. 1E). This method also identified the terminus of the megaplasmid at 187° opposite its *ori* and within the 25 kbp region generating abundant TtAgo-bound smDNAs. Cumulative GC skew analysis of the *T. thermophilus* HB8 strain similarly showed that the 15–18 nt DNA guides associated with TtAgo mapped opposite the origin of replication (fig. S1F).

Second, we used high-throughput, short-read DNA sequencing to identify *ori* and *ter*. In logarithmically growing bacteria, DNA replication typically initiates more often than it concludes. Consequently, chromosomal sequences from *ori* are overrepresented, while those from *ter* are under-represented in logarithmically growing cells relative to those that have reached stationary phase (*21, 22*). The ratio of logarithmic to stationary phase genomic sequencing coverage placed *ter* 183° clockwise from *ori* at 597,500 bp, again within the region producing abundant smDNAs bound to TtAgo (fig. S1G).

## Catalytically inactive TtAgo fails to accumulate high levels of smDNA guides

Guide acquisition by TtAgo is poorly understood. Purified TtAgo has been reported to cleave dsDNA without sequence specificity, suggesting that TtAgo itself initiates guide production (*23*). Supporting this idea, smDNAs do not co-purify with catalytically inactive TtAgo^D478A,D546A^ (henceforth <u>d</u>ouble point-<u>m</u>utant, TtAgo^DM^) when over-expressed in *E. coli* grown at 37°C (*5*). We produced both mutant and wild-type TtAgo in *E. coli* and purified each protein to apparent homogeneity. In agreement with earlier studies, we did not detect either smDNA-directed or sequence-independent cleavage of single-stranded target DNA during a 16 h incubation at 65°C, even when purified TtAgo^DM^, loaded with a smDNA guide, was present 45,000-fold above its *K_D_*, a 50-fold excess over the singlestranded DNA (ssDNA) target fully complementary to the DNA guide (fig. S2A). Under these same conditions, purified wild-type TtAgo, guided by the same smDNA, cleaved the ssDNA target within 1 h; ssDNA cleavage required the smDNA guide. However, we failed to detect any cleavage of dsDNA by wild-type, purified recombinant TtAgo, either in the presence or absence of a smDNA guide (fig. S2B), and TtAgo bound dsDNA 40-fold more weakly than ssDNA guides (fig. S2D). Our data suggest that TtAgo is a DNA-guided, single-stranded DNA endonuclease devoid of detectable sequence-independent double-stranded DNase activity.

Further evidence against a role of TtAgo in the initial generation of its own smDNA guides comes from analysis of the nucleic acids associated with TtAgo^DM^ immunoprecipitated from *T. thermophilus* grown at 65°C. As reported previously for TtAgo^DM^ expressed in *E. coli*, (*5*), in vivo, TtAgo^DM^ bound 25–35 nt RNA (Fig. 2A), and these small RNAs mapped throughout the genome. Unlike in *E. coli*, TtAgo^DM^ in vivo also bound smDNA guides that had the same length distribution as those associated with wild-type TtAgo, typically began with cytidine, and mapped to a region encompassing the chromosomal terminus (Fig. 2, B and C, and fig. S1D). Notably, 86% of the uniquely mapping smDNA sequences co-immunoprecipitated with TtAgo^DM^ also co-purified with TtAgo (Fig. 2D). However, the guides associated in vivo with the catalytically inactive protein were ~20 -fold less plentiful than for wild-type TtAgo, even though in vivo, wild-type and mutant TtAgo proteins accumulated to equivalent levels and were immunoprecipitated with the same efficiency; recombinant TtAgo and TtAgo^DM^ were both equally active and bound smDNA guides with similar affinities (Fig. 2E and fig. S2, C and D). We conclude that the TtAgo DNA endonuclease acts in smDNA loading or amplification, but not in their initial production.

**Fig. 2.**
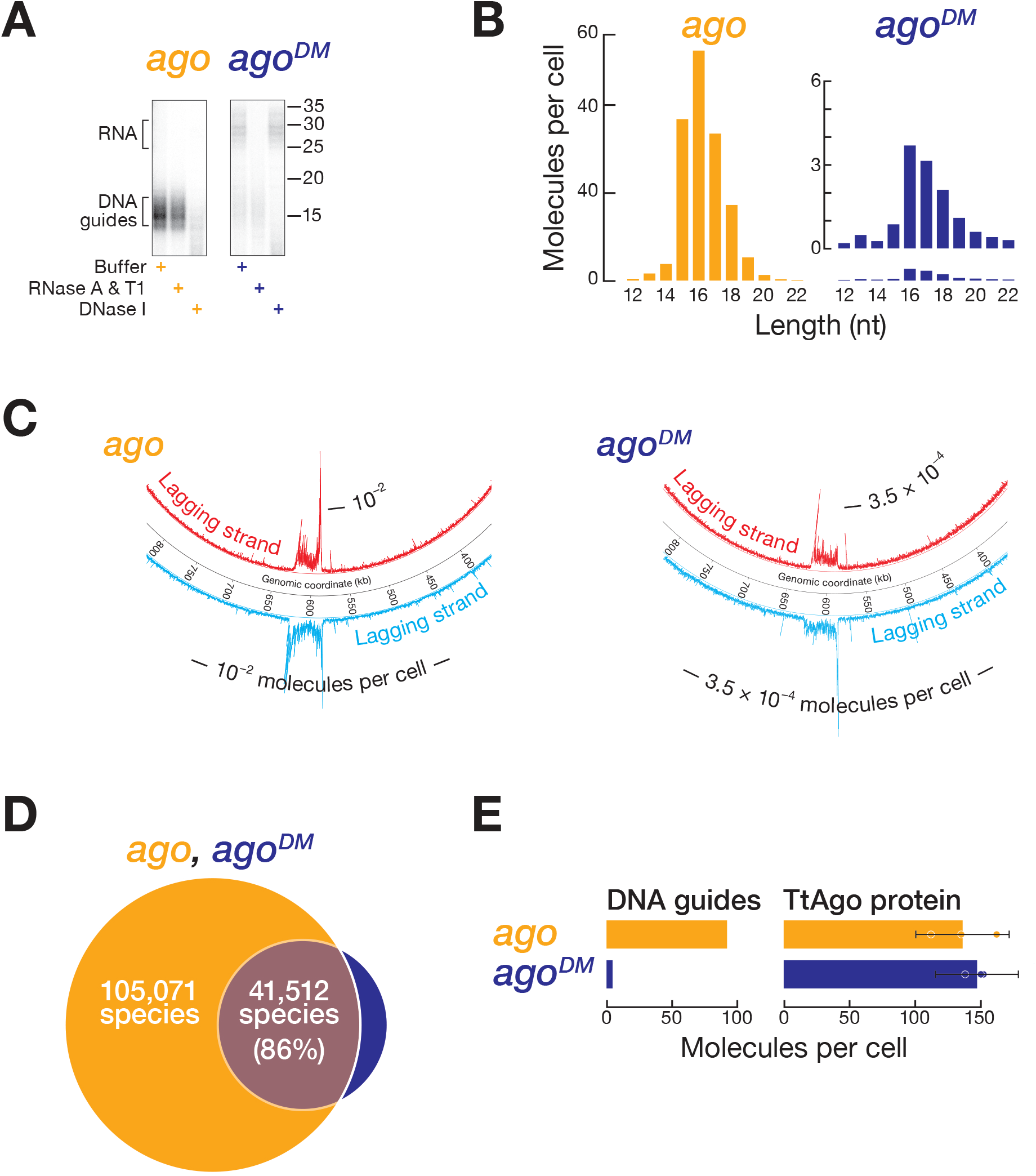
Comparison of smDNAs bound to TtAgo and TtAgo^DM^ in vivo. (**A**) Nucleic acids bound to TtAgo. (**B**) Length distribution of smDNA guides bound by TtAgo. (**C**) TtAgo-bound smDNA guides aligned to chromosomal terminus. (**D**) Comparison of guide sequences (≥ 10 ppm) bound to TtAgo. (**E**) Abundance of TtAgo proteins and associated smDNA guides in vivo. Data are from cells harvested at OD_600_ = 0.5.

## TtAgo and gyrase collaborate to terminate DNA replication

The observation that TtAgo binds smDNA guides from the chromosomal and megaplasmid termini suggests that in vivo the protein participates in DNA replication or genome maintenance. To explore this idea, we screened wild-type (*ago*) and null mutant (*Δago*) HB27 strains for their susceptibility to a panel of replication inhibitors and DNA damaging agents. We observed no difference in growth between *ago* and *Δago* grown in the presence of the DNA damaging agents 4-Nitroquinoline 1-oxide or methyl methanesulfonate; the DNA crosslinking agents cisplatin or mitomycin C; the ribonuclease reductase inhibitor hydroxyurea; or the gyrase subunit B inhibitor novobiocin. However, the *Δago* strain was more sensitive to the gyrase subunit A inhibitor ciprofloxacin than wild-type *ago* (Fig. 3A and fig. S3A). Increasing concentrations of ciprofloxacin slowed the growth of *Δago*, but they did not decrease the viability of the mutant strain, compared with *ago* (fig. S3, B and C). Under these conditions, the abundance of wild-type TtAgo-associated smDNA guides mapping to the terminus increased (fig. S4, A to C).

**Fig. 3.**
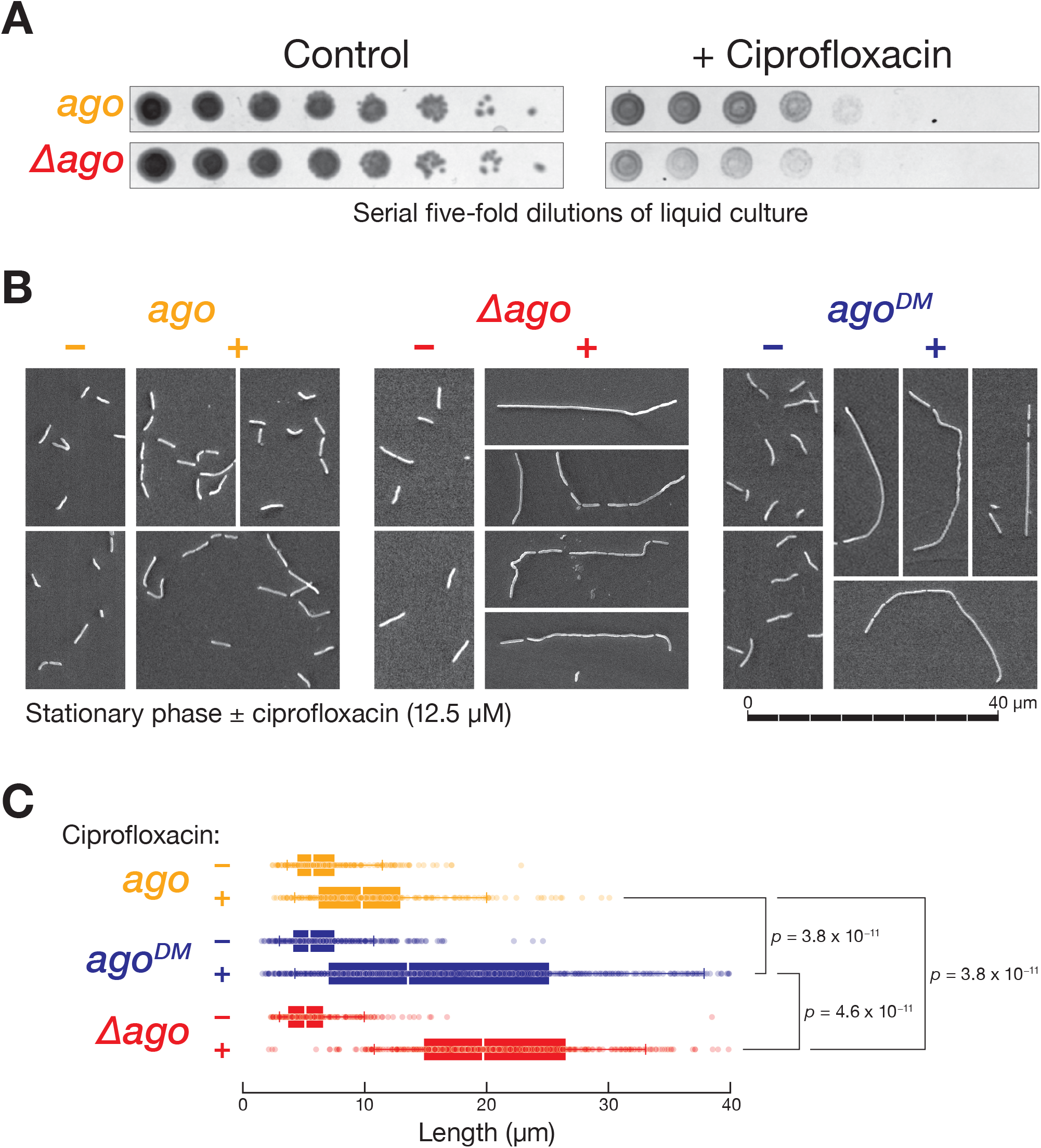
Effect of ciprofloxacin on *T. thermophilus* growth and morphology. (**A**) Growth susceptibility of *ago* and *Δago* to 13 μM ciprofloxacin. (B) Scanning electron microscopy analysis of strains grown in the presence or absence of 12.5 μM ciprofloxacin. (C) Length distribution of *T. thermophilus* cells grown in the presence or absence of 12.5 μM ciprofloxacin. Two-way ANOVA: ciprofloxacin and genotype effect on length (*F* (2, 2277) = 77.743, *p* < 2 × 10^−16^); ciprofloxacin effect on length (*F* (1, 2277) = 612.90, *p* < 2 × 10^−16^); genotype effect on cell length (*F* (2, 2277) = 3.537, *p* = 0.03). Tukey test: ciprofloxacin treated cells were significantly longer than untreated cells (7.1 μm, *p* = 3.8 x 10^−11^, 95% CI [6.5, 7.9]); treated *Dago* cells were longer than *ago* (2.8 μm, *p* = 3.8 × 10^−11^, 95% CI [2.8-3.4]; treated *ago^DM^* cells were longer than *ago* (1.7 μm, *p* = 3.8 × 10^−11^, 95% CI [1.5-2.0]); treated *Dago* cells were longer *ago^DM^* (1.6 μm, *p* = 4.6 × 10^−11^, 95% CI [1.3-1.9])

The mechanism by which the agents inhibit replication gives insight into the role of TtAgo in vivo. Hydroxyurea inhibits ribonucleotide reductase, depleting available dNTPs. The lack of differential susceptibility between *ago* and *Δago* suggests replication elongation does not rely upon TtAgo. The increased sensitivity of *Δago* to ciprofloxacin, which inhibits gyrase A, but not to novobiocin, which interferes with the B subunit of the gyrase heterodimer, further restricts the possible roles for TtAgo in DNA replication. Gyrase A cleaves both strands of dsDNA, forming a covalent DNA-enzyme bond. Gyrase B uses ATP energy to induce negative supercoiling (*24*). The negative twist generated by gyrase B is captured by re-ligation of the ends of the dsDNA by gyrase A. Gyrase can also use the same mechanism to unlink catenated dsDNA circles (*25, 26*). In *E. coli*, a second topoisomerase II enzyme, topo IV, decatenates the circular chromosomes generated by DNA replication. *T. thermophilus* lacks topo IV. Thus, the ability of gyrase A to break and rejoin dsDNA is predicted to decatenate replicated DNA in *T. thermophilus*. When gyrase A is inhibited by ciprofloxacin, *T. thermophilus* should therefore be unable to separate the catenated DNA circles generated by replication. Failure to decatenate daughter chromosomes is predicted to impair nucleoid segregation and prevent complete septation, causing individual cells to remain connected and form long filaments. Contrary to this expectation, wild-type *ago* grew essentially normally, showing only a modest increase in median cell length in the presence of ciprofloxacin (Fig. 3, B and C). How then does *T. thermophilus* complete replication without gyrase A function?

Successful nucleoid segregation and complete septation when gyrase A activity is inhibited requires TtAgo: both the *Δago* and *ago^DM^* strains formed long filaments when gyrase A was inhibited by ciprofloxacin (Fig. 3B). When the media contained ciprofloxacin, both *Dago* and *ago^DM^* cells were significantly longer than wild-type, TtAgo-expressing cells (*p* = 3.8 × 10^−11^ for both; Tukey test). Moreover, in the presence of ciprofloxacin, *Dago* cells were significantly longer than *ago^DM^* (*p* = 4.6 × 10^−11^; Tukey test), consistent with the small amount of DNA guides present in the catalytically inactive TtAgo^DM^ strain (Fig. 3C). We conclude that TtAgo, loaded with smDNA guides corresponding to the terminus, can facilitate completion of replication in *T. thermophilus*.

Transmission electron microscopy (TEM) and stimulated emission depletion (STED) microscopy images of wild-type and mutant *T. thermophilus* strains grown in ciprofloxacin underscore the requirement for TtAgo in nucleoid segregation and complete septation. By TEM, the daughter nucleoids of wild-type *ago* cells were set back from the cell wall (Fig. 4A). In contrast, the daughter nucleoids of *Dago* and *ago^DM^* cells lay close to the septal junctions within long filaments of incompletely separated cells. Longitudinal cross-sections of *Dago* and *ago^DM^* filaments showed that the nucleoids of adjacent segments were joined by a thin fiber that extended from cell to cell along the length of the filament. To determine whether this fiber corresponds to DNA, we stained the cells with PicoGreen, a dye whose fluorescent emission increases ~2,000-fold when bound to dsDNA (*27*), and examined them using STED microscopy (Fig. 4B). In the presence of ciprofloxacin, the thin fiber extending between cells stained brightly with PicoGreen in both *Dago* and *ago^DM^*, consistent with the cells being linked by dsDNA. No PicoGreen-staining material was observed between wild-type *ago* cells. We conclude that in the absence of both gyrase A and TtAgo function, newly replicated *T. thermophilus* chromosomal DNA cannot be decatenated, blocking completion of septation.

**Fig. 4.**
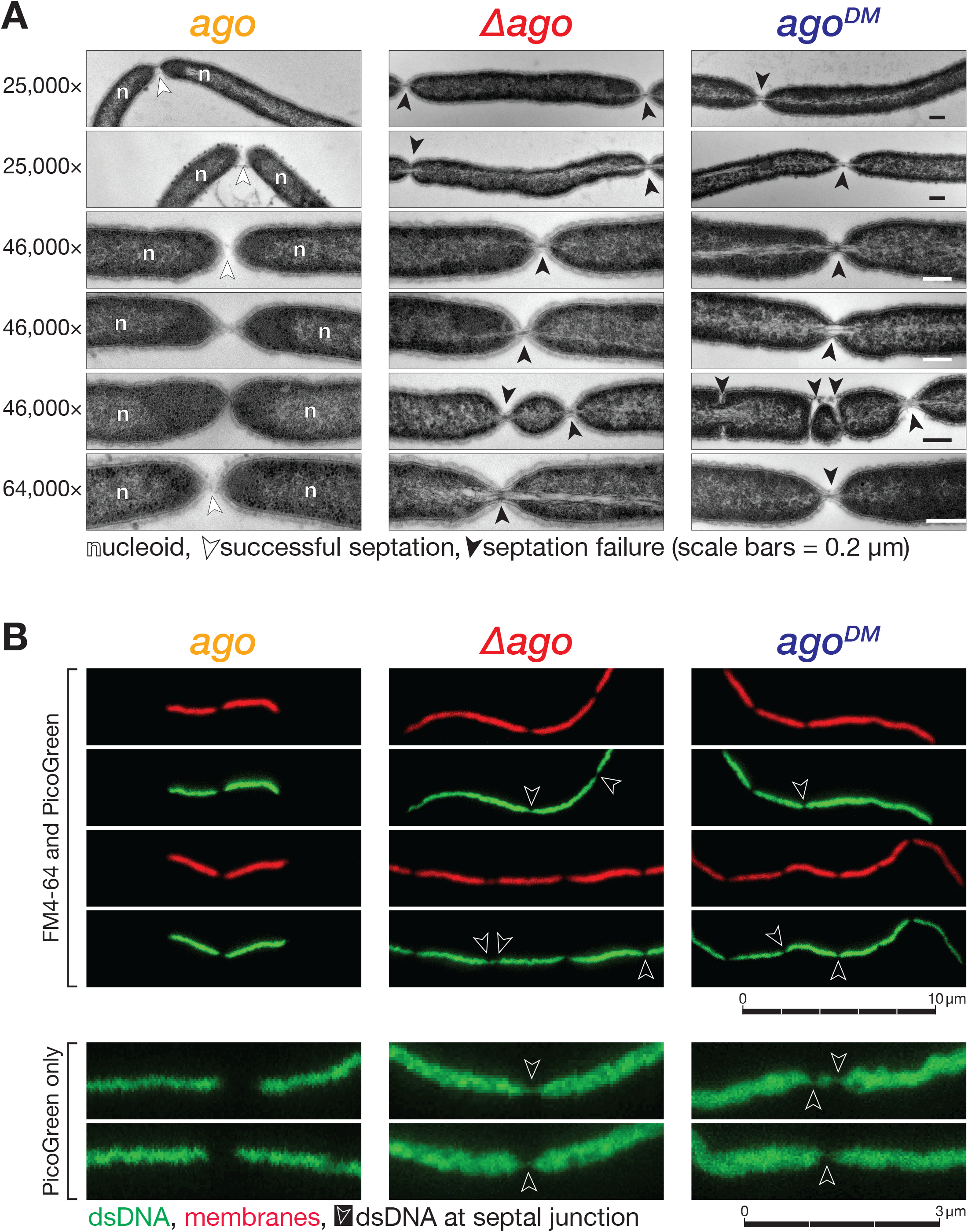
Effect of ciprofloxacin on nucleoid morphology. (**A**) Transmission electron microscopy cross-sectional analysis of *T. thermophilus* grown in the presence or absence of 12.5 μM ciprofloxacin. Multiple representative images are shown. (**B**) Stimulated emission depletion microscopy images of *T. thermophilus* grown in the presence or absence of 12.5 μM ciprofloxacin. PicoGreen detects dsDNA (green); FM4-64 detects membranes (red). Multiple dye images are two sets of identical cells and single dye images are independent, representative images.

## TtAgo binding partners implicate recombination in the completion of DNA replication

The identity of proteins associated with TtAgo suggests that the protein collaborates with recombination factors both to generate DNA guides and to separate catenated chromosomes (Fig. 5 and fig. S5A). The *T. thermophilus* genome encodes an AddA/AddB helicase-nuclease complex, which, like RecBCD, initiates repair of doublestrand breaks by homologous recombination and is required for completion of replication (*28–30*). Mass spectrometry analysis of the proteins that specifically coimmunoprecipitated with TtAgo identified both AddA and AddB, as well as the recombination factors Rad52 and ArgR (XerA), the ssDNA-binding protein SSB, and the histone-like protein HU, consistent with TtAgo acting at DNA. (TtAgo appears to bind sites of persistent ssDNA generally: RepA, which initiates DNA copying at *oriC*, also copurified with TtAgo, and TtAgo bound smDNAs mapping to the origin of replication [Fig. 1E]). Supporting the idea that TtAgo acquires its guides from the stalled replication forks that accumulate at the end of replication of circular chromosomes, TtAgo co-purified with PriA, which helps re-start stalled replication forks; RecJ, an exonuclease that repairs stalled forks; Topol, which relieves negative supercoiling behind replication forks; and PolA, which replaces the RNA primers on the lagging strand of the fork with DNA. Proteins acting at the end of replication (GyrA and GyrB), in DNA repair (UvrB), and in cytokinesis (FtsE), also co-immunoprecipitated with TtAgo. Many of these TtAgo interacting proteins likely bind via protein-protein interactions, because (1) their association persisted when the lysate was incubated with DNase before immunoprecipitation (Fig. 5 and fig. S5B), and (2) many of the proteins associated with wild-type TtAgo also associated with catalytically inactive TtAgo^DM^, despite the mutant protein binding >20-fold fewer smDNA guides (Fig. 5). Notably, the recombination proteins AddA and AddB, which co-purified with wild-type TtAgo even after DNase treatment, were not significantly associated with TtAgo^DM^, perhaps because they bind only TtAgo loaded with a smDNA guide. Figure S5C provides a complete list of proteins specifically co-immunoprecipitating with TtAgo.

**Fig. 5.**
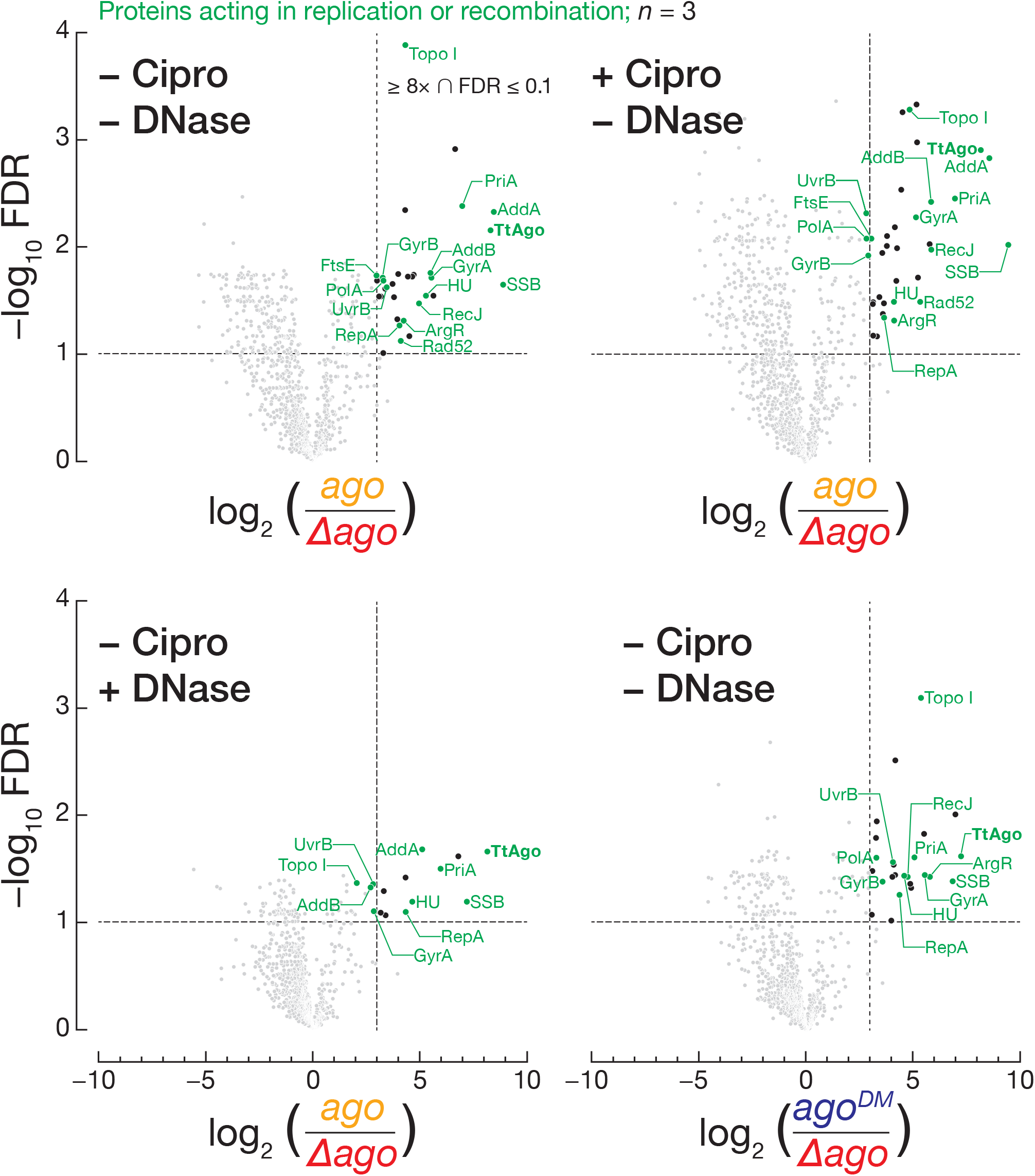
Identification of proteins associated with TtAgo. Proteins associated with TtAgo in *ago* grown 8 h in the absence or presence of 12.5 μM ciprofloxacin, compared to *Δago* null mutant at OD_600_ = 0.5 (top panels). Proteins specifically co-immunoprecipitating with TtAgo (*ago*) after DNase treatment or with TtAgo^DM^ (*ago^DM^*) at OD_600_ = 0.5 compared to *Δago* null mutant (bottom panels). Dashed lines: FDR = 0.1 (horizontal) and 8× enrichment (vertical). Green: proteins known to function in replication or recombination. Black: other proteins; Grey: proteins with FDR > 0.1 and/or < 8× enrichment.

## Discussion

Argonaute proteins defend against viral infection, silence transposons, inhibit transformation by plasmids, direct transcriptional silencing, promote mRNA decay, and repress translation. Our data expand the list of Argonaute functions to include ensuring successful replication of circular chromosomes: in vivo, TtAgo, binds ~16 nt DNA guides derived from the terminus of replication and facilitates decatenation, enabling subsequent nucleoid segregation and cytokinesis. In the laboratory, we can detect the function of TtAgo in chromosomal replication only when GyrA is inhibited; future experiments will be needed to determine whether TtAgo confers a long-term selective advantage to *T. thermophilus* under more natural conditions.

How does TtAgo, a DNA-guided, DNA-binding, DNA endonuclease facilitate chromosome decatenation? Under normal physiological conditions, the gyrase heterodimers first cut both DNA strands of one newly replicated chromosome in the terminus, forming a covalent DNA-enzyme complex with one strand of each of the two resulting gate-segments (G-segments) (*24, 31*) (Fig. 6A). The other daughter chromosome, the transfer-segment, is passed through this break, decatenating the two circles. Finally, the G-segments covalently linked to Gyrase A are ligated to each other, regenerating a circular chromosome and allowing the two chromosome copies to partition between daughter cells. In the presence of ciprofloxacin, the G-segments are trapped as DNA-enzyme intermediates, blocking decatenation and eliciting filamentation. We propose that TtAgo-guide complex can bypass this block by binding complementary ssDNA sequences in the terminus (*32*). Once bound, TtAgo resolves catenanes either (1) by slicing the ssDNA to generate a double-strand break which can then be resected by TtAgo-associated AddAB, triggering recombination or (2) by simply bringing recombination factors, such as Rad52, ArgR, or AddAB, to the terminus. We favor the second model, because the severity of filamentation in the catalytically inactive *ago^DM^* strain was significantly less than in *Δago*. Our data suggest that smDNA guides derived from the terminus direct TtAgo to the terminus, and that TtAgo associated proteins, rather than endonucleolytic cleavage of DNA, drive decatenation itself. Published observations provide additional support for this model: (1) expression of the TtAgo MID-PIWI domain in *E. coli* enhances sequence-directed recombination (*33*), and (2) in *E. coli*, a region identical in size and location to the *T. thermophilus* smDNA locus is a site of hyperrecombination (*34, 35*).

**Fig. 6.**
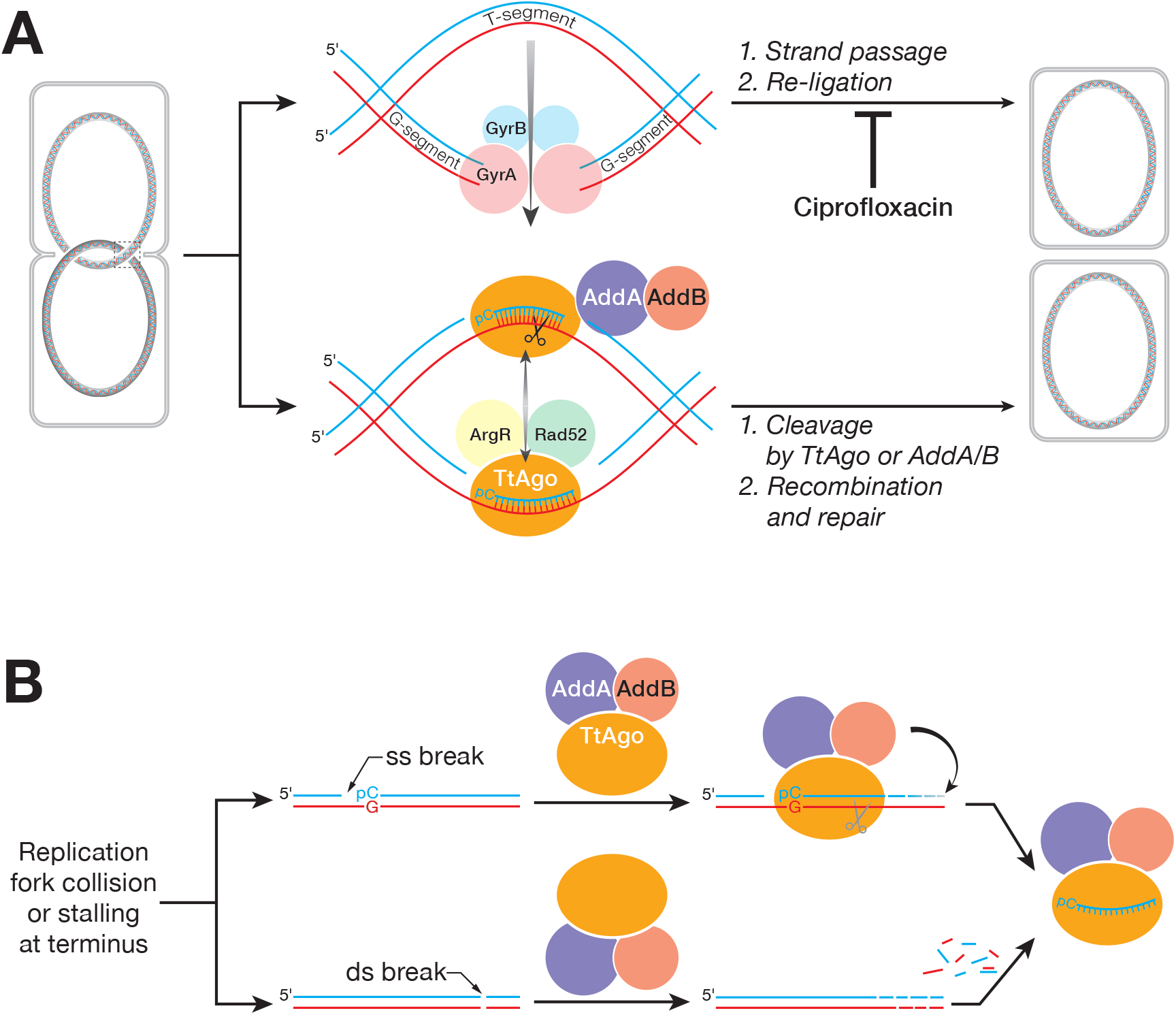
Models of TtAgo function in vivo. Proposed models for (**A**) how Gyrase and TtAgo act to resolve catenated daughter chromosomes, and (**B**) how TtAgo might acquire DNA guides.

How does TtAgo acquire guides from the terminus region? As replication forks approach the terminus, forks stall, often causing nicks and dsDNA breaks (*36, 37*) (Fig. 6B). We can imagine two mechanisms by which TtAgo can exploit DNA breaks to obtain guides. First, TtAgo may bind the free 5’ monophosphate at nicks in dsDNA, perhaps at nicks beginning with a 5’ cytidine, reflecting the protein’s preference for targets bearing a guanosine across from the first guide nucleotide (*23, 38*). Although TtAgo prefers to bind ssDNA, the protein’s in vivo concentration (>200 nM) far exceeds its in vitro *K_D_* (30 nM) for binding 5’ monophosphorylated dsDNA. Next, TtAgo would cleave the target strand, providing an entry point for its associated AddAB complex to degrade both the target DNA strand and trim the 3’ end of the guide strand, liberating a ~16 nt ssDNA guide bound to TtAgo. Alternatively, AddAB may generate free smDNAs that its associated TtAgo can then bind. The finding that TtAgo^DM^ is largely devoid of guides favors the first model, but a detailed dissection of guide acquisition may require development of a cell-free *T. thermophilus* lysate that recapitulates the process.

Do Argonaute proteins decatenate chromosomes or otherwise participate in DNA replication, recombination, or repair in other prokaryotes? Although Argonaute proteins can be found across prokaryotes, only some are predicted to retain cleavage activity (*1*). Our data suggest that species harboring active Argonaute proteins comprise the most promising candidates for exploration. A broad survey of these strains may require the development of a metagenomics approach to identifying Argonaute guides in complex mixtures of bacteria. Given the growing problem of bacterial resistance to antibiotics, identifying Argonaute functions in prokaryotes could provide a starting point for developing drugs that block DNA replication or cytokinesis in combination with existing antibiotics.

## ACKNOWLEDGEMENTS

We thank Christina Baer for help with imaging; Daniel Tom and Douglas Matthews of Leica for assistance with STED microscopy; Nicholas Rhind for critical analysis and guidance; Ahmet Ozturk for Circos software training; and members of the Zamore laboratory for discussions and comments on the manuscript.

## Author contributions

Conceptualization: P.D.Z. and S.M.J.; methodology: S.M.J., I.G., G.M.H. and A.D.; performed experiments: S.M.J., H.Z., K.J., L.S. and A.D.; provided resources: G.M.H. and B.U.; formal analyses: I.G., S.M.J., K.J., A.D.; writing – original draft: S.M.J.; supervision and funding acquisition: P.D.Z.

## Data and materials availability

Raw sequencing data is available from XXX using accession number YYY. Mass spectrometry data is available from XXX using accession number YYY.

## Methods summary

Detailed materials and methods can be found in the supplementary materials. Briefly, we prepared marked wild-type and mutant strains by homologous recombination using vectors containing a thermostable kanamycin resistance gene for selection, and the genomes verified by long-read sequencing. Small DNA guides bound to TtAgo were prepared for sequencing using a splint ligation technique (*39*) and visualized using Circos software. Proteins co-immunoprecipitating with TtAgo were identified by LC-MS/MS. Spot plating assays were performed by plating serial dilutions of logarithmic phase *T. thermophilus* on gellan-gum fortified agar containing the inhibitor of interest. Gross external morphology was imaged by SEM. Lengths of filaments were determined from DIC microscopy images of fixed cells. The nucleoid was characterized by TEM and STED microscopy.

## REFERENCES AND NOTES

1. S. Ryazansky, A. Kulbachinskiy, A. A. Aravin, MBio 9, (2018).

2. L. Lisitskaya, A. A. Aravin, A. Kulbachinskiy, Nat Commun 9, 5165 (2018).

3. D. C. Swarts et al., Nat Struct Mol Biol 21, 743 (2014).

4. K. S. Makarova, Y. I. Wolf, J. van der Oost, E. V. Koonin, Biol Direct 4, 29 (2009).

5. D. C. Swarts et al., Nature 507, 258 (2014).

6. D. C. Swarts et al., Nucleic Acids Res 43, 5120 (2015).

7. A. Zander, P. Holzmeister, D. Klose, P. Tinnefeld, D. Grohmann, RNA Biol 11, 45 (2014).

8. T. Oshima, K. Imahori, Int J Syst Bacteriol 24, 102 (1974).

9. R. A. D. Williams, K. E. Smith, S. G. Welch, J. Micallef, R. J. Sharp, International Journal of Systematic and Evolutionary Microbiology 45, 495 (1995).

10. A. Henne et al., Nat Biotechnol 22, 547 (2004).

11. H. Li, G3 (Bethesda) 9, 1249 (2019).

12. N. Ohtani, M. Tomita, M. Itaya, J Bacteriol 192, 5499 (2010).

13. M. Gao et al., Cell Res 24, 532 (2014).

14. C. Oliver, J. L. Santos, M. Pradillo, Front Plant Sci 5, 177 (2014).

15. W. Wei et al., Cell 149, 101 (2012).

16. K. M. Michalik, R. Bottcher, K. Forstemann, Nucleic Acids Res 40, 9596 (2012).

17. I. Schmidts, R. Böttcher, M. Mirkovic-Hösle, K. Förstemann, Nucleic Acids Res 44, 8261 (2016).

18. A. Levy et al., Nature 520, 505 (2015).

19. A. Grigoriev, Nucleic Acids Res 26, 2286 (1998).

20. S. Schaper et al., J Mol Biol 299, 655 (2000).

21. M. Hawkins, S. Malla, M. J. Blythe, C. A. Nieduszynski, T. Allers, Nature 503, 544 (2013).

22. A. Koren, I. Soifer, N. Barkai, Genome Res 20, 781 (2010).

23. D. C. Swarts et al., Mol Cell 65, 985 (2017).

24. J. M. Berger, S. J. Gamblin, S. C. Harrison, J. C. Wang, Nature 379, 225 (1996).

25. A. Aubry, L. M. Fisher, V. Jarlier, E. Cambau, Biochem Biophys Res Commun 348, 158 (2006).

26. A. W. Debowski et al., PLoS One 7, e33310 (2012).

27. V. L. Singer, L. J. Jones, S. T. Yue, R. P. Haugland, Anal Biochem 249, 228 (1997).

28. G. A. Cromie, J Bacteriol 191, 5076 (2009).

29. J. Kooistra, G. Venema, J Bacteriol 173, 3644 (1991).

30. J. Courcelle, B. M. Wendel, D. D. Livingstone, C. T. Courcelle, DNA Repair (Amst) 32, 86 (2015).

31. A. Sugino, C. L. Peebles, K. N. Kreuzer, N. R. Cozzarelli, Proc Natl Acad Sci U S A 74, 4767 (1977).

32. P. Nurse, C. Levine, H. Hassing, K. J. Marians, J Biol Chem 278, 8653 (2003).

33. L. Fu et al., Nucleic Acids Res 47, 3568 (2019).

34. T. M. Hill, Escherichia coli and Salmonella typhimurium: Cellular and Molecular Biology (Neidhardt, FC, ed.), 2nd edit 1602 (1996).

35. J. Louarn, F. Cornet, V. François, J. Patte, J. M. Louarn, J Bacteriol 176, 7524 (1994).

36. A. Kuzminov, Proc Natl Acad Sci U S A 98, 8241 (2001).

37. B. Michel et al., Proc Natl Acad Sci U S A 98, 8181 (2001).

38. C. S. Smith et al., Nat Commun 10, 272 (2019).

39. I. Olovnikov, K. Chan, R. Sachidanandam, D. K. Newman, A. A. Aravin, Mol Cell 51, 594 (2013).

