## Supplemental Materials for "A DNA-guided Argonaute Protein Functions in DNA Replication in *Thermus thermophilus*"

**This PDF file includes:**

Materials and Methods
Figs. S1 to S5
Table S1
References

### Materials and Methods

#### Strains and mutants

Studies used wild-type *Thermus thermophilus* HB27 (ATCC BAA-163, DSM 7039) or HB8 (ATCC 27634), wild-type incorporating a thermostable kanamycin resistance gene (*ago*::*htk*), or incorporating the resistance marker in the context of catalytically inactive mutant TtAgo (D478A,D546A, referred to as $ago^{DM}$::*htk* or $ago^{DM}$) or deletion of the entire TtAgo locus (*Δago::htk* or *Δago*). Mutants were obtained by constructing vectors and recombining into HB27 or HB27 mutants. Vectors were constructed by Gibson assembly (*1*) and inserted into the Sma1 site of pUC19.*htk* was amplified from plasmid RDB3436 (Riken Data Bank, Japan), *hyg* was amplified from pTRH1T (*2*), adding 300 bp of upstream and downstream sequence from *T. thermophilus* genome; $TtAgo^{DM}$ was amplified from pWUR703 (*3*). Homologous recombination was performed as described (*4*) with selection on 200 µg/mL kanamycin A TR-plates or 40 µg/mL hygromycin B TR-plates. *Δago*::*htk* and *ago*::*htk* strains were prepared by recombining into the wild type HB27 strain. $ago^{D478A,D546A}$::*htk* was prepared by first recombining $ago^{D478A,D546A}$::*hyg* into the *Δago*::*htk* strain and subsequent replacing *hyg* with *htk*. Genomic sequencing of all strains confirmed that mutations or deletions existed as designed and in all copies of the genome.

#### *Thermus thermophilus cultures*

Strains were grown in ATCC-697 media (HB27; 0.8% (w/v) polypeptone, 0.4% (w/v) yeast extract, 0.2% (w/v) sodium chloride, pH 7.5 with NaOH) or TB media (HB8; 0.8% (w/v) polypeptone, 0.4% (w/v) yeast extract, 0.3% (w/v) sodium chloride, 0.5 mM calcium chloride, 1 mM magnesium chloride, pH 7.5 with NaOH). For HB27 strains, overnight cultures were started by adding glycerol stocks to 5 mL ATCC-697 media and incubating at 65°C with 300 rpm shaking for 12–15 h. Experimental cultures were started by seeding fresh ATCC-697 media with overnight culture to achieve 2–3 × $10^7$ cells/mL, then incubating at 65°C with 250 rpm. shaking at 250 rpm. For growth in ciprofloxacin, 40 mL of a 100 mL seed culture at $OD_{600}$ = 0.5 was added to a flask containing 166 µg (5 × $10^{-4}$ moles) ciprofloxacin (17850, Sigma, USA) dried onto the bottom from an 0.05 M stock in 0.1 N HCl, achieving a final concentration of 12.5 µM ciprofloxacin. Cultures were allowed to grow at 65°C for 8 h with 250 rpm shaking achieving $OD_{600}$ = 2–2.5.

Bacteria were plated on TR-plates (1.5% (w/v) gellan gum [Alfa Aesar J63423A1], 0.4% (w/v) polypeptone, 0.2% (w/v) yeast extract, 0.1% (w/v) sodium chloride, 1.5 mM calcium chloride, 1.5 mM magnesium chloride, pH 7.5 with NaOH) were prepared, dried overnight and stored at 4°C until use. Following inoculation, plates were incubated at 65°C for 18–24 h in plastic bags containing damp paper towels. For spot plating assays, the agar was equilibrated to 70°C before adding the inhibitor. Each strain was grown to $OD_{600}$ = 0.5 and serially diluted fivefold. 250 µL in a 96-well plate. A frogger was dipped into the well and stamped onto an agar plate containing inhibitor. Between agar plates, the frogger was dipped into a 96-well plate containing fresh media that was changed before use with a different inhibitor. Plates were incubated at 65°C for 15 min right side up, then place in sealed bags containing damp paper towels and incubated upside down at 65°C for 18 h.

Colony diameter and colony-forming unit measurements

A liquid culture grown to $OD_{600}$ = 0.5 was diluted 1:500,000, and then 25 µL was spread on TR-plates containing vehicle (0.1N HCl), 7.5 µM, 10 µM, 12.5 µM or 15 µM ciprofloxacin. Plates were incubated at 65°C for 15 min right side up, then were placed in sealed bags containing damp paper towels and incubated upside down at 65°C for 24 h. Images were acquired using an Amersham Imager 600 (GE healthcare, USA) and colony diameter determined using Image J (Fuji). The number of colonies was counted without respect to colony size.

Growth curves

An overnight cultures was used to seed with $5 \times 10^8$ cells into 40 mL of growth media containing 166 μg ($5 \times 10^{-4}$ moles) ciprofloxacin as described above. The culture was sampled hourly for the first 12 hours and once at 14 h and 24 h. Culture samples were diluted 1:4 to measure $OD_{600}$.

Filamentation assay

Cultures were grown in the absence or presence of 12.5 μM ciprofloxacin as described above. After 8 h, 80 μL of the culture was added to 320 μL fresh media, allowed to cool to room temperature, and then 1 mL 4% paraformaldehyde in Dulbecco's phosphate buffered saline (PBS, 137 mM NaCl, 2.7 mM KCl, 8 mM $Na_2HPO_4$, 2 mM $KH_2PO_4$; Gibco, 14190-144) was added and incubated 10 min at room temperature. The suspension was centrifuged at 1,000 × *g* at 4°C for 5 min and the supernatant removed. Cells were washed twice with 1 mL PBS. The final cell pellet was re-suspended in 150 μL PBS, and then 5 μL of the suspension was placed on a thin 1% (w/v) agar pad on a microscope slide. ProLong Gold (Invitrogen, P369931) mounting agent was added to the agar pad, capped with a cover slip, and the slides were allowed to cure overnight at room temperature. Slides were imaged on a Leica DMi8 by DIC microscopy with a 63×/NA 1.4 oil immersion objective. Fields with minimally overlapping bacteria were identified and stitched images of 3×3 panels acquired. Bacterial lengths were measured in Image J, counting only bacteria with distinct boundaries. Data was log-transformed and subject to a test of normality (Shapiro-Wilk test) and a test of homoscedasticity (Levene's test) before proceeding to two-way analysis of variance (ANOVA) to explore for significant independent and interacting effects of genotype and ciprofloxacin treatment, followed by post-hoc testing with Tukey's honestly significant difference test for pairwise comparisons.

Lysate preparation

Bacteria were grown to logarithmic ($OD_{600}$ = 0.5) or stationary phase (8 h after $OD_{600}$ = 0.5), collected at 7,000 × *g* for 10 min, and washed three times with 1× Dulbecco′s PBS. Cells were re-suspended in lysis buffer (25 mM HEPES pH 7.5, 100 mM potassium chloride, 5 mM magnesium chloride, 1 mM dithiothreitol (DTT), 1 mM 4-(2-aminoethyl)benzenesulfonyl fluoride hydrochloride, 0.3 µM aprotinin, 40 µM Bestatin, 10 µM E-64, 10 µM leupeptin hemisulfate) to make a 10% (w/v) suspension and lysed (1) using a microfluidizer (Microfluidics M-110P), passing the suspension once at 18,000 psi with cooling or (2) by sonication (Fisher Scientific, Sonic Dismembrator FB-120) on ice for four 30 s cycles at 40% amplitude with 30 s rest between cycles. Lysates were cleared by centrifugation at 20,000 × *g* for 20 min at 4°C, aliquoted into tubes, flash frozen in liquid nitrogen, and stored at −80°C.

Purification of recombinant TtAgo and $TtAgo^{DM}$

Wild-type or mutant TtAgo ($TtAgo^{D478A,D546A}$ or $TtAgo^{DM}$) cloned and expressed from pET-SUMO (Invitrogen K30001) in *E. coli* BL21-DE3 was purified as described (*5*) except: (1) After cleavage and removal of the 6×His-SUMO tag, wild-type or mutant TtAgo was additionally purified by HiTrap SP HP (GE Healthcare) chromatography, dialyzed against 3 × 2 L 20 mM HEPES-KOH pH 7.4, 250 mM potassium acetate, 3 mM magnesium acetate, 0.1 mM 2,2′,2″,2‴-(ethane-1,2-diyldinitrilo)tetraacetic acid, 10% glycerol (w/v), 5 mM DTT (TtAgo) or 30 mM HEPES-KOH, pH 7.4, 250 mM potassium acetate, 1 mM DTT, 0.01% Igepal CA-630, 20% [v/v] glycerol ($TtAgo^{DM}$). Purified protein was aliquoted into tubes, flash frozen in liquid nitrogen, and stored at −80°C. Protein was quantified by Bradford Assay using BSA as standard or by amino acid analysis (Protein Structure Core Facility, University of Nebraska Medical Center).

Anti-TtAgo antibody

Untagged, recombinant TtAgo (1.3 mg/animal) was excised from a 10% SDS-PAGE gel and submitted for development of rabbit anti-TtAgo polyclonal antibodies (Pocono Rabbit Farm and Lab, Inc., Pennsylvania). After 70 days, rabbits were exsanguinated, and antibodies purified from sera using recombinant TtAgo covalently attached to cyanogen bromide-activated Sepharose (Sigma Aldrich, C9142). Column was washed with 20 mM sodium phosphate (pH 7.2) and eluted with 0.1 M glycine (pH 2.8) immediately neutralized with 1 M Tris-HCl (pH 9.0). Antibody-containing fractions were combined, aliquoted, flash frozen, and stored at −80°C.

Filter binding assays

Binding assays were performed as described (*6*) except that for stoichiometric binding to compare TtAgo and $TtAgo^{DM}$, protein was incubated with ssDNA guides at 75°C for 15 min and then allowed to cool to room temperature before assaying and 10 nM radiolabeled guide was used

Western blotting

Samples were adjusted to 60 mM Tris-HCl pH 6.8, 2% (w/v) sodium dodecyl sulfate, 10% (w/v) glycerol, 0.1 M DTT, 0.1% (w/v) bromophenol blue, heated at 95°C for 5 min, and resolved by electrophoresis through a 4–20% Tris-glycine gel (Invitrogen, XP04025). Proteins were transferred at 40V for 14 h (Bio-Rad Mini Trans-Blot) from the gel to PVDF membrane (Millipore, IPVH00010), blocked with blocking buffer (Rockland Immunochemicals, MB-070) for 30 min at room temperature, and incubated for 60 min at room temperature with affinity-purified rabbit anti-TtAgo polyclonal antibody diluted

1:4,000 in blocking buffer. Membrane was washed for 10 min at room temperature with TBST (20 mM Tris-HCl pH 7.6, 150 mM sodium chloride, 0.01% (w/v) Tween-20) five times, incubated with goat anti-rabbit IRDye 800CW secondary antibody (LI-Cor Biosciences, 926-32211, 1:15,000 in blocking buffer) at room temperature for 60 min protected from light, washed four times for 10 min each at room temperature with TBST, and once with TBST containing 0.01% (w/v) sodium dodecyl sulfate. Buffer was replaced with TBST, and membrane imaged (Odyssey Infrared Imaging System, Li-Cor).

Immunoprecipitation

Affinity-purified anti-TtAgo rabbit polyclonal antibody was bound to Protein A paramagnetic beads (Invitrogen, 10002D) for 15 min, rotating end-over-end. Antibody was crosslinked to beads only for mass spectrometry experiments and was performed using bis(sulfosuccinimidyl)suberate (Invitrogen, 21580) according to the manufacturer's protocol; for all other immunoprecipitations no crosslinking of antibody was performed. Crosslinked antibody-bead complexes were used at 8 μg antibody/mg total protein; non-crosslinked antibody-bead complexes were used at 4 μg antibody/mg total protein. Antibody-bead complexes were incubated with cell lysates for 60 min at room temperature and washed three times with lysis buffer and the proteins were eluted in 25 mM Tris-HCl pH 7.5, 150 mM sodium chloride, 1% (w/v) Igepal CA-630, 1% (w/v) sodium deoxycholate, 2% (w/v) sodium dodecyl sulfate, 1 mM DTT or in 120 mM Tris-HCl pH 6.8, 4% (w/v) sodium dodecyl sulfate, 20% (w/v) glycerol, 0.2 M DTT and 0.2% bromophenol blue (w/v) with heating at 95°C for 5 min.

Nucleic acid guide characterization

Following immunoprecipitation of TtAgo from *T. thermophilus* lysate (2.5 mg total protein), immune complexes on beads were suspended in 200 mM Tris-HCl pH 7.5, 300 mM sodium chloride, 25 mM 2,2′,2″,2‴-(Ethane-1,2-diyldinitrilo)tetraacetic acid, 2% (w/v) sodium dodecyl sulfate (400 µL). Proteinase K (Denville Scientific, CB3200-7) was added (f.c. 200 µg/mL). The suspension was heated at 37°C for 3 h, and complete protein digestion confirmed by SDS-PAGE with silver staining (Pierce, 24612). Nucleic acids were isolated by ethanol precipitation, and taken up in 50 mM Bis-tris-propane-HCl pH 6, 1 mM magnesium chloride, 0.1 mM zinc chloride and treated (1) with Antarctic phosphatase (NEB, M0289) or (2) without enzyme (to identify 5′-phosphorylation state) at 37°C for 30 min. Phosphatase was inactivated at 80°C for 2 min. The phosphatase treated nucleic acids were forward labeled with γ-$^{32}$P-ATP in 70 mM Tris-HCl pH 7.6, 10 mM magnesium chloride, 5 mM DTT, and 15 U T4 Polynucleotide kinase (NEB, M0201)at 37°C for 30 min. The non-phosphatase treated nucleic acids were exchange labeled with the above buffer supplemented with 1 mM ADP. Free nucleotides were removed with a MicroSpin G-25 column (GE Healthcare). A portion of the flow through was treated with RQ1 DNase (2U, Promega, M6101) or RNase A/T1 cocktail (0.5U/20U, Invitrogen, AM2286) in 40 mM Tris-HCl pH 8.0, 10 mM magnesium sulfate, 1 mM calcium chloride at 37°C for 30 min. Reactions were stopped by adding an equal volume of 200 mM Tris-HCl pH 7.5, 300 mM sodium chloride, 25 mM 2,2′,2″,2‴-(Ethane-1,2-diyldinitrilo)tetraacetic acid, 2% (w/v) sodium dodecyl sulfate containing Proteinase K (f.c. 200 µg/mL) and incubated at 50°C for 15 min. The reaction was extracted with 25:24:1 phenol:chloroform:isoamyl alcohol pH 8.0, precipitated with ethanol, and resolved by 15% urea polyacrylamide gel electrophoresis. The gel was dried and imaged with a Typhoon FLA 7000 phosphorimager (GE Healthcare).

Small DNA sequencing

Immunoprecipitates on antibody beads were incubated in 400 µL of 200 mM Tris-HCl pH 7.5, 300 mM sodium chloride, 25 mM 2,2′,2″,2‴-(Ethane-1,2-diyldinitrilo)tetraacetic acid, 2% (w/v) sodium dodecyl sulfate containing Proteinase K (f.c. 200 µg/mL) at 37°C for 3 h. Nucleic acids were recovered by ethanol precipitation, treated with RNase A/T1 (0.5 U and 20 U, respectively; Invitrogen, AM2286) in 50 mM Tris-HCl pH 7.9, 100 mM sodium chloride, 10 mM magnesium chloride, 1 mM DTT at 37°C for 30 min, then extracted with 25:24:1 phenol:chloroform:isoamyl alcohol (pH 8.0), and the DNA was precipitated with ethanol and dissolved in water (20 µL). To one quarter of the immunoprecipitated DNA (5 µL) was added a total of either 3 or 12 fmoles of an equimolar mixture of six distinct oligonucleotide spike-ins (Table S1) in 5 µL. Next 10 µL formamide loading buffer (98% (w/v) deionized formamide, 10 mM EDTA, 0.025% (w/v) xylene cyanol, 0.025% (w/v) bromophenol blue) was added, the sample heated at 95°C for 5 min and then resolved by electrophoresis through a denaturing 15% polyacrylamide gel. Separate gels were used to purify DNA from each strain to prevent cross-contamination. The region of the gel corresponding to 12–25 nt was excised, crushed and incubated in 1.2 mL 30 mM HEPES-KOH pH 7.4, 300 mM sodium chloride, 25 mM EDTA, 2% (w/v) sodium dodecyl sulfate at room temperature for 14 h, and following filtration of the gel fragments. DNA was recovered by ethanol precipitation and dissolved in water (20 µL). Before ligation, duplexes of adapter and bridge oligonucleotides were annealed by adding 192 µL 100 µM bridge oligonucleotide and 160 µL 100 µM of the corresponding adapter oligonucleotide (Table S1), 40 µL buffer (500 mM Tris-HCl, pH 7.5, 100 mM magnesium chloride, 10 mM rATP, 100 mM DTT) to 8 µL water and heated to 95˚C for 2 min, and then allowed to cool to room temperature. To the size-selected DNA in water (10 µL) was added 2 µL buffer (500 mM Tris-HCl, pH 7.5, 100 mM magnesium chloride, 10 mM rATP, 100 mM DTT), 2.5 µL pre-annealed 3′-adapter duplex (100 pmoles; Table S1), 2.5 µL pre-annealed 5′-adapter duplex (100 pmoles; Table S1), 2 µL PEG8000 (50% v/v in water) and 1 µL T4 DNA Ligase (400 U, NEB M0202), and the reaction heated at 30˚C for 4 h. The reaction was adjusted to 300 mM sodium chloride, precipitated with ethanol and purified by electrophoresis through a denaturing 10% polyacrylamide gel. Gels were stained by SYBR-Gold, excised, and bands excised and eluted into 0.3 M sodium chloride, and finally recovered by precipitation with ethanol. The library was amplified and purified as reported for small RNA sequencing (*7*).

Long DNA sequencing

Immunoprecipitated DNA was isolated as above for small DNA sequencing with the exception that no size selection step was used. The DNA average length by Bioanalyzer (Agilent, USA) was 1,156 bp. The sequencing library was prepared by sequentially performing end-repair, A-tailing, Y-shaped adaptor ligation, and PCR amplification as described (*8*) and sequenced using a NextSeq 500 (Illumina) to obtain 75 nt, paired-end reads.

Library preparation for origin and terminus sequencing

Cultures were grown as above with the exception that cell pellets were isolated at $OD_{650}$ 0.1 for logarithmic phase and $OD_{650}$ 1.4 for stationary phase (*9*). Genomic DNA was isolated as described above for genomic sequencing, and the DNA was sheared (E220 Evolution, Covaris Inc., USA) for 210 s with duty cycle 10%, intensity 5 and 180 cycles/burst to achieve ~250 bp fragments. The sequencing library was prepared by sequentially performing end-repair, A-tailing, Y-shaped adaptor ligation, and PCR amplification as described (*8*) with the exception that unique molecular identifiers were used as described (*10*). The libraries were sequenced using a NextSeq 500 (Illumina) to obtain 75 nt, paired-end reads.

Genomic sequencing

Stationary phase *Thermus thermophilus* cells (0.4 g) were collected by centrifugation and resuspended in 20 mM Tris-HCl, pH 7.5, 300 mM sodium chloride, 25 mM 2,2′,2″,2‴-(Ethane-1,2-diyldinitrilo)tetraacetic acid, 2% (w/v) sodium dodecyl sulfate.

Genomic DNA was isolated as described (*11*) except that the DNA was not sheared during isolation. Genomic DNA was submitted to the University of Massachusetts Medical School Deep Sequencing Pac-Bio Core for library preparation, sequencing using a PacBio RS or Sequel II, and genome assembly using HGAP2 or HGAP4.

Analysis of high-throughput sequencing data

For small DNAs, sequences were filtered (Phred quality score ≥20 for all nucleotides), and the 3′ adapter sequence (5′-TGGAATTCTCGGGTGCCAAGG-3′) removed. Sequences of spike-in synthetic DNAs were identified allowing no mismatches. Sequences of small DNAs ≥14 nt in length were aligned onto our de novo genome assembly allowing no mismatches. For multiply mapping sequences, the number of read was divided by the number of locations to which the small DNA mapped (i.e., multi-mapping reads were apportioned). For origin and terminus mapping, PCR duplicates were removed using unique molecular identifiers (*10*), and total coverage per 1,000 bp for the chromosome was determined for each growth phase. The ratio of logarithmic:stationary phases of identical windows was calculated and smoothed using the Loess smoothed option in Igor Pro (WaveMetrics, USA) using a smoothing factor = 1 and a quadratic regression order (i.e., 2).

Stimulated emission depletion microscopy

Cultures were grown in the presence of 12.5 µM ciprofloxacin as described above. After 8 h, 100 µL was diluted into 400 µL ice-cold Hank's balanced salt solution (HBSS; $Na_2PO_4$, NaCl, $NaHCO_3$, $KH_2PO_4$, KCl; pH 7.4; mOsm = 290; Invitrogen, 14175). A 1 mL volume of 4% (v/v) paraformaldehyde in HBSS was added and incubated on ice for 10 min. The suspension was centrifuged at 1,000 × *g* at 4°C for 3 min. The supernatant was removed, and the pellet re-suspended in 1 mL HBSS. The cells were centrifuged as before, the supernatant removed, and then the cells were re-suspended in 1 mL

HBSS. Of this suspension, 150 µL were diluted into 600 µL HBSS to make a dilute cell suspension. To 100 µL of the dilute cell suspension was added 100 µL PicoGreen (Invitrogen, P7581) diluted 1:200 in TE (10 mM Tris-HCl pH 7.5, 1 mM EDTA), which was then incubated at room temperature in the dark for 10 min. Stained cells (10 µL) were added to a #1.5 cover slip (VWR, 48393172) and allowed to air dry in the dark for 60 min. To the dried cells was added ProLong Glass mounting agent (Invitrogen, P36982), a slide was placed on the cover slip and the slide was allowed to cure for 15 h prior to imaging. For samples stained sequentially with FM4-64fx (Invitrogen, F34653) before and PicoGreen, the protocol was as above except that after initial dilution and prior to fixation, 2.5 µL FM4-64fx (1000 µg/mL in water) was added to 500 µL diluted culture to a final concentration of 5 µg/mL, incubated on ice for 1 min in the dark, and the fixation was performed as above except it was carried out in the dark. Staining with PicoGreen was then carried out as above. Images were acquired on a Leica SP8 STED with a 100×/NA 1.40 oil immersion objective. PicoGreen was excited at 488 nm wavelength and STED was performed at using 592 nm using 80% laser strength for depletion and filter range of 496-583 nm. Only confocal images were acquired (no STED) with FM4-64, which was excited at 514 nm and a filter range of 628-699 nm. A Z-projection of maximal intensity was assembled for each channel using ImageJ.

Scanning electron microscopy

Cultures were grown with or without ciprofloxacin as described above. Either 4 mL (for cells at $OD_{600}$ = 0.5) or 2 mL (for cells grown an additional 8 h) were added without cooling to six-well plates (3.5 cm; Corning, 3516) containing poly-L-lysine-coated coverslips and incubated for 5 min at room temperature. For 4 mL samples, 2 mL media was removed. Next, 2 mL pre-warmed (55°C) fixation buffer (2.5% glutaraldehyde (v/v) in 0.1 M sodium cacodylate buffer (pH 7.2)) was added to each well and incubated 10 min at room temperature. The media was then replaced with 2 mL fixation buffer, and the cells incubated for 1 h at room temperature. Samples were rinsed three times with fixation buffer, dehydrated using a graded series of ethanol in 20% increments, ending with 100% ethanol, followed by three washes with 100% ethanol and then critical point dried with AutoSAMDRI-815 (Tousimis, USA) for 30 min. Cover slips were mounted on aluminum SEM stubs using double-sided carbon tape (8 mm; SPI, 05072-AB) , grounded with colloidal silver paint, sputter coated with 12 nm gold-palladium (60:40; Ted Pella Inc., 91111), and imaged using secondary electron (SEI) mode using a Quanta 200 FEG MKII scanning electron microscope (ThermoFisher, USA).

<u>Transmission electron microscopy</u>

Cultures were grown with or without ciprofloxacin as described above. Either 10 mL (for cells at $OD_{600}$ = 0.5) or 5 mL (for cells grown an additional 8 h) were added without cooling to six-well plates (3.5 cm; Corning, 3516) containing poly-L-lysine-coated coverslips and incubated for 5 min at room temperature. For 10 mL samples, 5 mL media was removed. Next, 5 mL pre-warmed (55°C) fixation buffer (2.5% glutaraldehyde (v/v) in 0.1 M sodium cacodylate buffer, pH 7.2). The media was then replaced with 2 mL fixation buffer, and the cells incubated for 1 h at room temperature. Samples were rinsed three times with fixation buffer, dehydrated using a graded series of ethanol in 20% increments, followed by three changes of 100% ethanol. Samples were infiltrated with a mixture of 1:1 absolute ethanol:SPI-Pon 812 epoxy resin (SPI, 02660-AB) overnight. After five changes of fresh SPI-Pon 812 epoxy resin for 1 h each at room temperature, samples were embedded in place in the wells in fresh epoxy resin for 48 h at 68°C. Epoxy wells were isolated with a jeweler's saw and submerged in liquid nitrogen until the plastic bottom separated from the resin. Resin was trimmed and glued sample side up to a blank Pon 812 epoxy resin block and sectioned (~70 nm) with an ultramicrotome (Leica UC7, Germany) using a diamond knife. Sections were mounted on copper support grids and incubated sequentially with lead citrate

(Reynold's formulation) for 3 min at room temperature and uranyl acetate (4% (v/v) in 25% (v/v) ethanol) for 4 min at room temperature. Sections were examined by at 100KV using a Philips CM10 transmission electron microscopy (Field Electron and Ion Co., USA) equipped with a charge-coupled device camera (ES1000W Erlangshen, Gatan Inc., USA).

Cleavage assays

*Cleavage comparison of TtAgo and TtAgo$^{DM}$*. Reactions (30 µL) were performed in final concentrations of 20 mM HEPES-KOH pH 7.4, 125 mM potassium acetate, 0.01% (w/v) Igepal CA-630 (Sigma), 5 mM dithiothreitol and 100 nM 5′-$^{32}$P-radiolabeled target and variable concentrations of $MnCl_2$, guide, TtAgo or TtAgo$^{DM}$ were added to complete the reaction. The mixture was heated at 65°C. Aliquots (7 µL) were removed at 0, 1 and 16 h, quenched into 100 µL Proteinase K buffer (200 mM Tris-HCl pH 7.5, 300 mM NaCl, 25 mM EDTA, 2% (w/v) sodium dodecyl sulfate) and flash frozen in liquid nitrogen then stored at -20˚C. Aliquots were thawed, 100 µL Proteinase K (0.2 mg/mL f.c.) in Proteinase K buffer was added and the mixture was heated at 50°C 1 h. The reaction was extracted with 1 volume of 24:25:1 phenol:chloroform:isoamyl alcohol (pH 8.0) and precipitated with ethanol. The samples were taken up in 25 µL formamide loading buffer (98% (w/v) deionized formamide, 10 mM EDTA (pH 8.0), 0.025% (w/v) xylene cyanol, 0.025% (w/v) bromophenol blue), heated at 95°C for 5 min and then resolved by electrophoresis through a denaturing 15% polyacrylamide gel. Imaged using a Typhoon FLA 7000IR phosphorimager (GE Healthcare).

*Cleavage of AT- or GC-rich targets*. Targets were identical to those described by Swarts et al. (*12*). Oligonucleotides were 5′ radiolabeled with γ-$^{32}$P-ATP (Perkin Elmer, NEG035C005MC) using T4 polynucleotide kinase (NEB, M201) following the manufacturer′s instructions. Free ATP was removed using a MicroSpin G-25 column

(GE Healthcare), and the oligonucleotides were further purified by denaturing 7.5% polyacrylamide gel electrophoresis. The 98 bp radiolabeled double-strand DNA substrates were prepared using PCR amplification of a 98 nt ssDNA template with a 5′-radiolabeled primer (prepared as above) and an unlabeled primer to achieve a substrate with the radiolabel incorporated on the strand complementary to the ssDNA guide. PCR was performed using Phusion polymerase (NEB, M0530), desalted (Qiagen, 28106), then purified by native 7.5% (29:1 acrylamide:bis-acrylamide) polyacrylamide gel electrophoresis. DNA was recovered by incubating the gel slice in XYZ for X h at Y°C, and then concentrated by ethanol precipitation. Cleavage assays were as described (*12*). Briefly, target substrates (20 nM ssDNA or 8 nM dsDNA) were incubated with TtAgo (50 nM) with or without ssDNA guide (60 nM) in 10 mM Tris-HCl pH 8.0, 0.5 mM manganese chloride, 250 mM sodium chloride for 16 h at 65°C. Reactions were stopped by adding 100 mM Tris-HCl pH 7.5, 150 mM sodium chloride, 12.5 mM 2,2′,2″,2‴-(ethane-1,2-diyldinitrilo)tetraacetic acid, 1% (w/v) sodium dodecyl sulfate, 200 ng/µL Proteinase K) and incubating at 65°C for 30 min., extracted with 25:24:1 phenol:chloroform:isoamyl alcohol (pH 8.0), precipitated with ethanol, and resolved by denaturing 7.5% polyacrylamide gel electrophoresis, and imaged using a Typhoon FLA 7000IR phosphorimager (GE Healthcare). Decade marker (Invitrogen, AM7778) labeled with γ-$^{32}$P-ATP were used as size standards.

Mass spectrometry

*Sample preparation*. Immunoprecipitation of TtAgo from lysate prepared by sonication using antibody covalently crosslinked to the beads was as described above except that after incubation with lysate, the antibody-beads were washed once with lysis buffer and then were re-suspended in 100 µL Turbo DNase buffer and 4U (2 µL) Turbo DNase (Ambion AM2238) and incubated at 37°C for 30 min. Water was used instead of DNase in the control incubation. Beads were washed three times with lysis buffer and eluted with 25 mM Tris-HCl pH 7.5, 150 mM sodium chloride, 1% (w/v) Igepal CA-630, 1% (w/v) sodium deoxycholate, 2% (w/v) sodium dodecyl sulfate, 1 mM DTT at 95°C for 5 min. Immunoprecipitates were reduced with (30 µL 0.2 M per 5 mg sample) for one h at 55°C, alkylated with iodoacetamide (30 µL 0.5 M per 5 mg sample) for 45 min in the dark at room temperature. Each sample was loaded onto S-Trap microcolumns (Protifi, USA) according to the manufacturer's instructions. Briefly, 3 µL of 12% phosphoric acid and 165 µL of binding buffer (90% methanol, 100 mM triethylammonium bicarbonate [TEAB]) were added to each sample. Samples were loaded onto spin columns and centrifuged at 4,000 × *g* for 30 s. After three washes with binding buffer, 20 µL containing 500 ng of trypsin in 50 mM TEAB was added to the microcolumn and incubated at 47°C for one h. Peptides were eluted using 40% acetonitrile in 0.5% acetic acid, followed by 80% acetonitrile in 0.5% acetic acid, and concentrated in a SpeedVac.

*LC-MS/MS analysis*. Each sample (1 µg) was loaded onto a 75 µm × 2 cm, C18, 3 µm, 100 Å PepMap 100 pre-column (Acclaim, Thermo Scientific) in series with a 50 m × 75 µm ID, 2 µm, 100 Å PepMap RSLC C18 EASY-Spray column analytical column ( ThermoFisher Scientific) using the autosampler of an Easy nLC 1000 (Thermo Scientific).. Peptides were eluted into the Orbitrap QExactive HF-X mass spectrometer (ThermoFisher Scientific) using a 5.9–22.28% gradient of acetonitrile in 0.5% acetic acid in 120 min, a 22.28–33.2% gradient of acetonitrile in 0.5% acetic acid in 20 min, and finally a 33.2–80 gradient of acetonitrile in 0.5% acetic acid in 10 min. The full scan was acquired with a resolution of 60,000 (@ *m*/*z* 200), a target value of $3 \times 10^6$ and a maximum ion time of 45 ms. Following each full MS scan, twenty data-dependent MS/MS spectra were acquired. The MS/MS spectra were collected with a resolution of 15,000, an AGC target of $1 \times 10^5$, a maximum ion time of 120 ms, one microscan, 2 m/z isolation window, fixed first mass of 150 m/z, dynamic exclusion of 30 sec, and Normalized Collision Energy (NCE) = 27.

*Data analysis*. MS/MS spectra were searched against the *T. thermophilus* reference proteome using Andromeda (*13*). Protein quantification between samples was performed using the summed eXtracted Ion Current (XIC) of all isotopic clusters within MaxQuant 1.5.2.8 (*13*). Mass tolerance was set to 10 ppm for MS1 and MS2 searches. False discovery rate (FDR) filtration was performed first on the peptide level and then on the protein level. Both filtrations were done at 1% FDR using a standard target-decoy database approach. Proteins identified with less than two unique peptides were excluded from analysis. Bioinformatics analysis was performed with Perseus and Microsoft Excel. Benjamini-Hochberg corrected Student's t-test was used to calculate FDR to identify proteins specifically associated with TtAgo.

**Data Availability:** strain sequencing, smDNA-seq, smRNA-seq, mass spectra

**Supplementary References**


1. D. G. Gibson *et al.*, *Nat Methods* **6**, 343 (2009).
2. A. Fujita, Y. Misumi, Y. Koyama, *Plasmid* **67**, 272 (2012).
3. D. C. Swarts *et al.*, *Nature* **507**, 258 (2014).
4. Y. Hashimoto, T. Yano, S. Kuramitsu, H. Kagamiyama, *FEBS Lett* **506**, 231 (2001).
5. Y. Wang *et al.*, *Nature* **456**, 921 (2008).
6. L. M. Wee, C. F. Flores-Jasso, W. E. Salomon, P. D. Zamore, *Cell* **151**, 1055 (2012).
7. B. W. Han, W. Wang, C. Li, Z. Weng, P. D. Zamore, *Science* **348**, 817 (2015).
8. Z. Zhang, W. E. Theurkauf, Z. Weng, P. D. Zamore, *Silence* **3**, 9 (2012).
9. M. Hawkins, S. Malla, M. J. Blythe, C. A. Nieduszynski, T. Allers, *Nature* **503**, 544 (2013).
10. Y. Fu, P. H. Wu, T. Beane, P. D. Zamore, Z. Weng, *BMC Genomics* **19**, 531 (2018).
11. M. R. Green, J. Sambrook, *Cold Spring Harb Protoc* **2017**, (2017).
12. D. C. Swarts *et al.*, *Mol Cell* **65**, 985 (2017).
13. J. Cox *et al.*, *J Proteome Res* **10**, 1794 (2011).

**Fig. S1. Characterization of TtAgo expression and bound nucleic acids.** (**A**) Full immunoblot from Fig 1A. Full immunoblot from Fig. 1A (outlined areas) (**B**) Full immunoblot of Fig. 1B (outlined areas). (**C**) Long DNAs bound to TtAgo. (**D**) Position-weight matrices of smDNAs bound to TtAgo or $TtAgo^{DM}$ (**E**) GC-content of various genomic regions. For ± 100 nt genomic context, all small DNAs were aligned by their 5′-end prior to analysis. (**F**) Small DNAs associated with TtAgo in vivo in strain HB8, aligned to the chromosome and megaplasmid. (**G**) DNA content from HB27 genomic sequencing calculated as logarithmic/stationary phase ratios in 1000 bp windows of summed coverage; line represents Loess smoothing.

Jolly et al., Figure S1

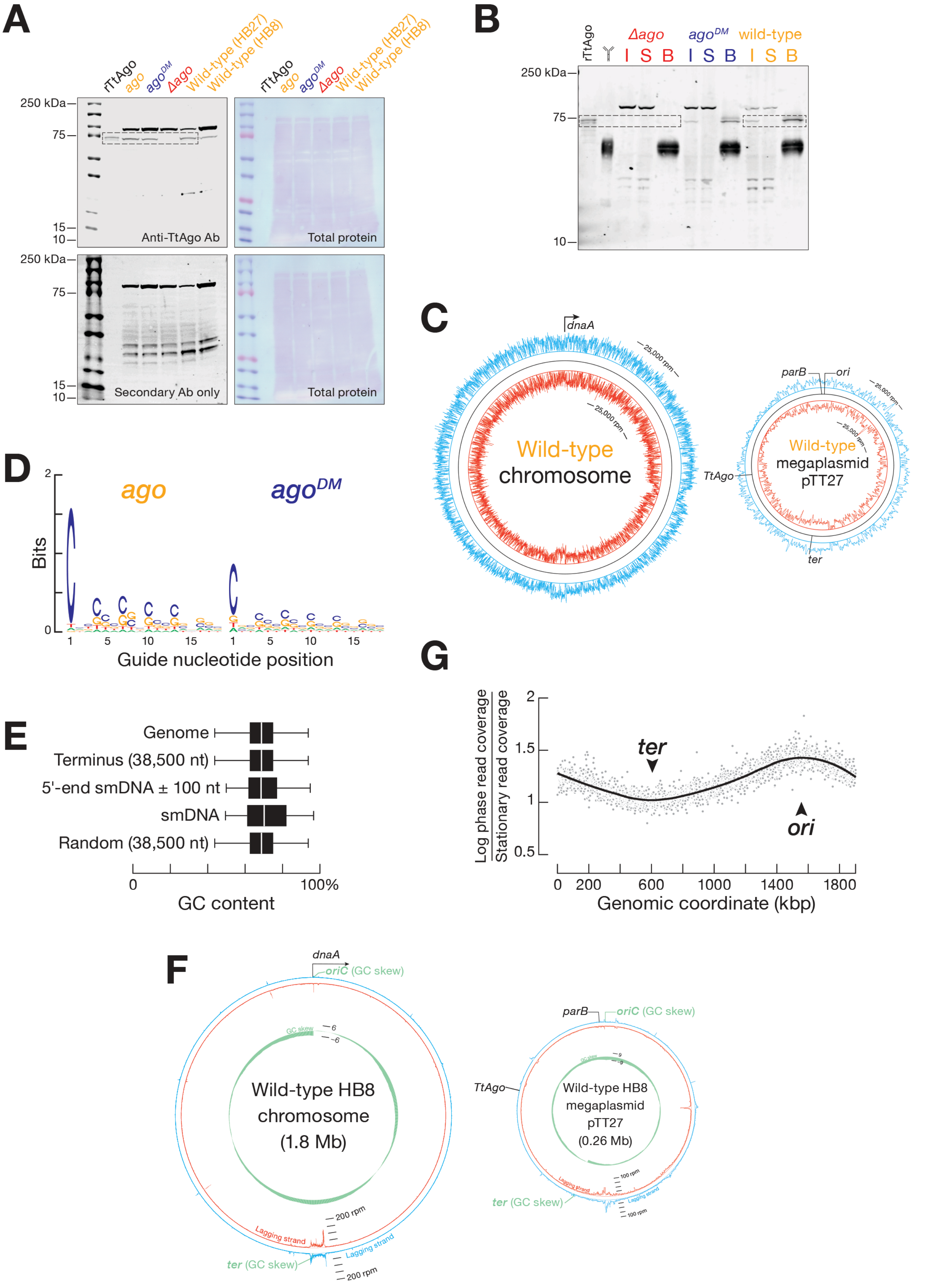

**Fig. S2 Characterization of TtAgo and TtAgo$^{DM}$.** (**A**) ssDNA target cleavage assays using TtAgo and TtAgo$^{DM}$. (**B**) TtAgo cleavage of ssDNA or dsDNA targets with high or low GC content. (**C**) Immunoblots of immunoprecipitations of TtAgo and TtAgo$^{DM}$ with and without 12.5 μM ciprofloxacin at logarithmic or stationary (8 h after adding ciprofloxacin dosing) phases. (**D**) Measured affinities of TtAgo and TtAgo$^{DM}$ various nucleic acids.

Jolly et al., Figure S2

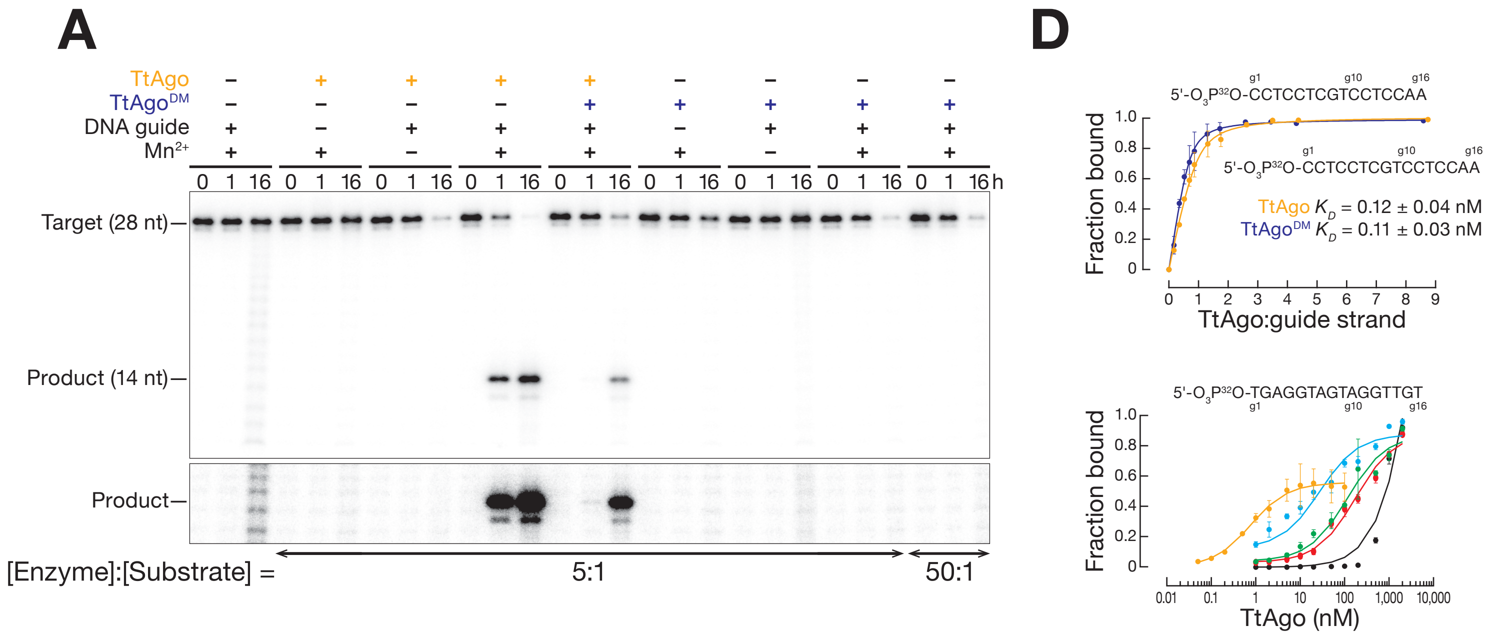


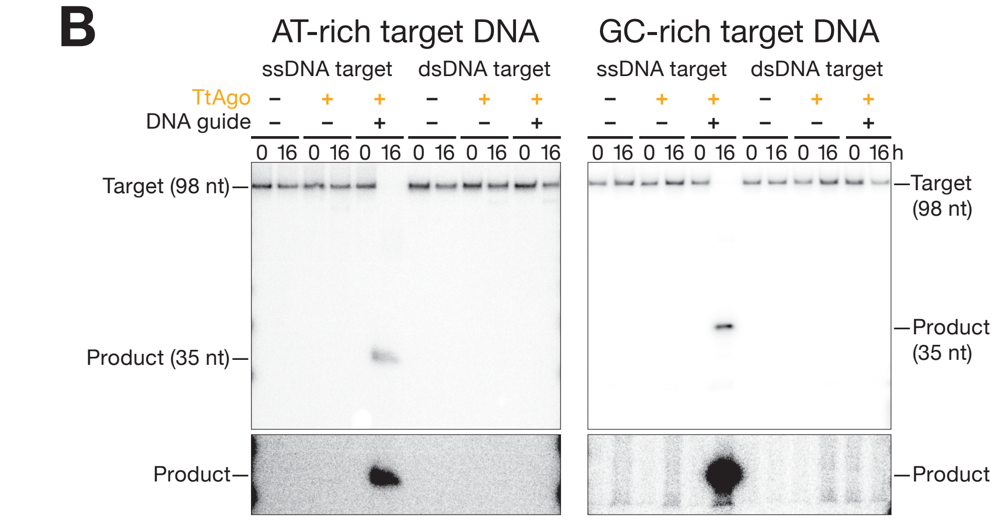


| Guide | $K_D$ (nM) | $K_{Relative}$ |
|---|---|---|
| ssDNA | 0.7 ± 0.3 | 1 |
| dsDNA | 30 ± 20 | 40 |
| DNA:RNA | 120 ± 50 | 170 |
| ssRNA | 180 ± 30 | 260 |
| dsRNA | 6,000 ± 1,000 | 8,600 |

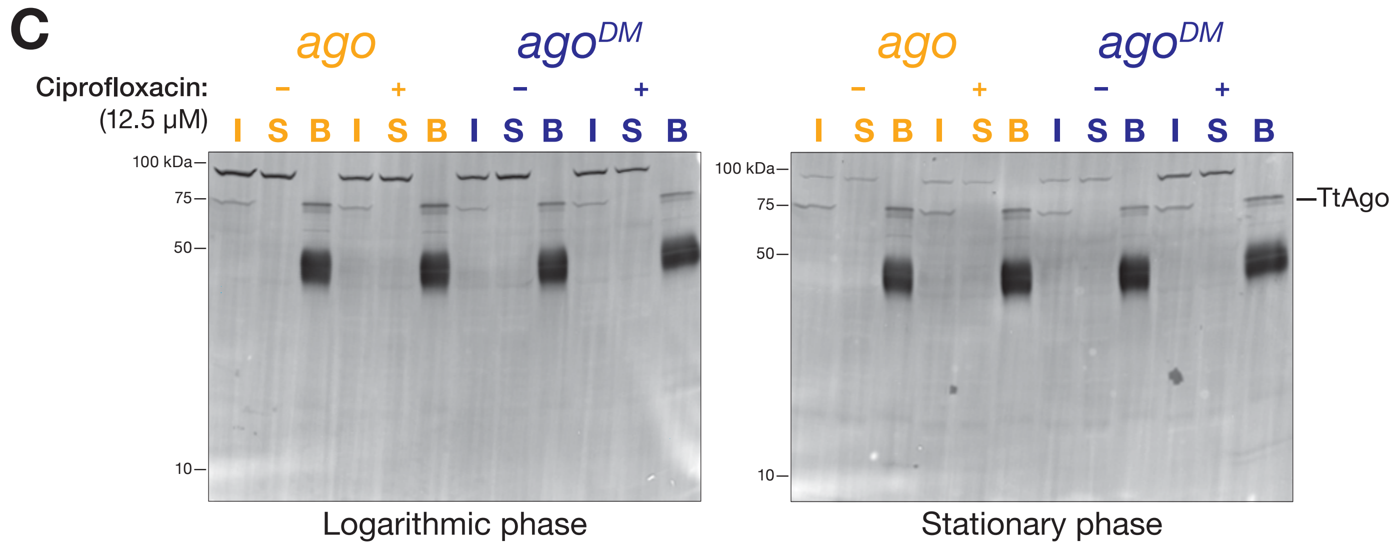

**Fig. S3. Growth of *T. thermophilus* HB27 strains in the presence of inhibitors.** (**A**) Spot assay of bacteria 5-fold serially diluted on plates containing the highest concentration at which bacterial growth was detected for each inhibitor. (**B**) Colony diameter of kanamycin-marked strains grown in the presence of different ciprofloxacin concentrations. (**C**) Growth kinetics of kanamycin-marked strains in the presence of 12.5 µM ciprofloxacin.

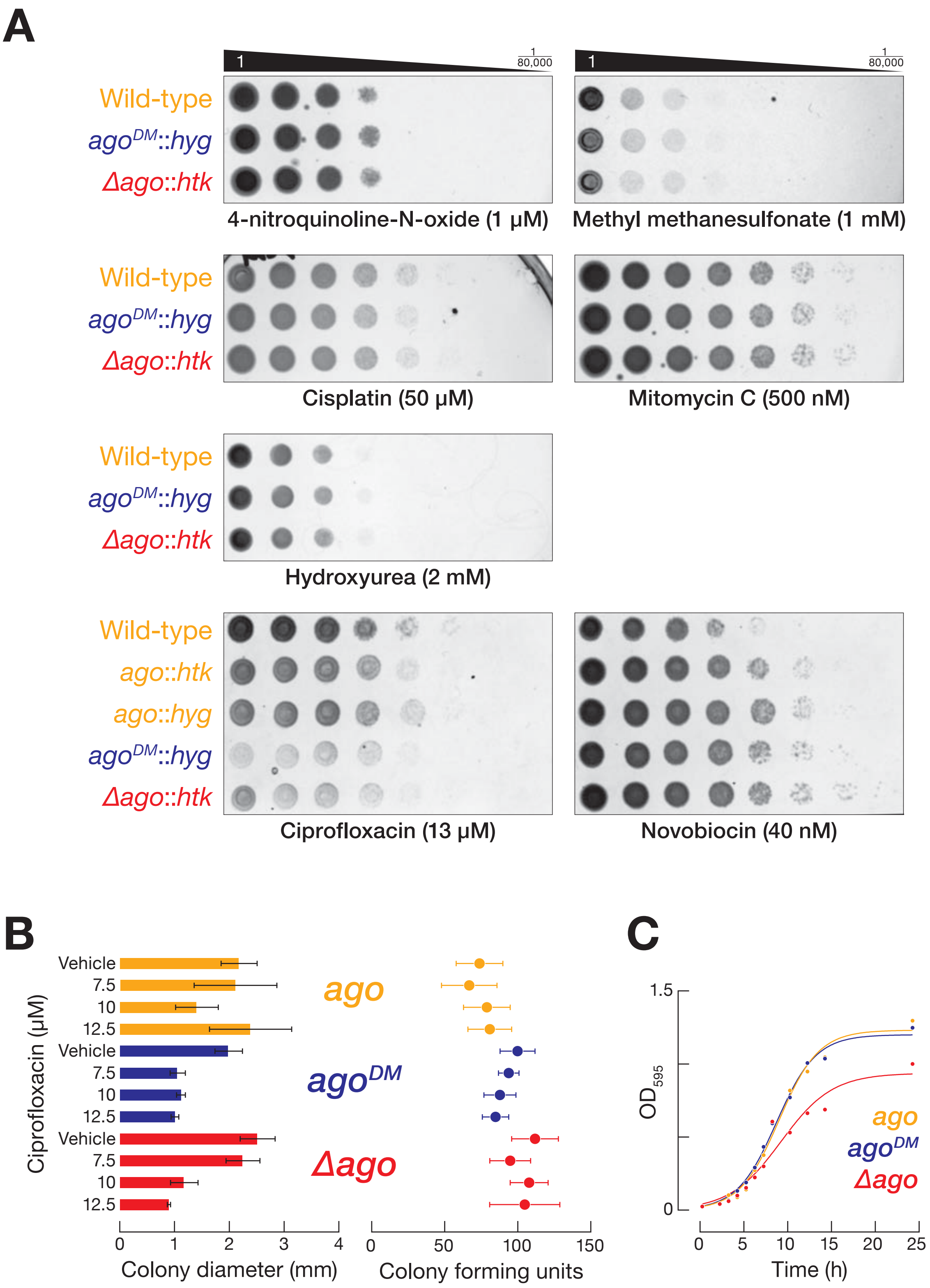
A
1
1/80,000
Wild-type
agoDM::hyg
Δago::htk
4-nitroquinoline-N-oxide (1 μM)
Methyl methanesulfonate (1 mM)
Cisplatin (50 μM)
Mitomycin C (500 nM)
Hydroxyurea (2 mM)
ago::htk
ago::hyg
Ciprofloxacin (13 μM)
Novobiocin (40 nM)
B
Ciprofloxacin (μM)
Vehicle
7.5
10
12.5
ago
agoDM
Δago
Colony diameter (mm)
Colony forming units
C
OD595
Time (h)
1.5
0

**Fig. S4. Characterization of *ago* and *ago$^{DM}$* grown in the absence or presence of 12.5 μM ciprofloxacin for 8 h; t = 0 h represents cultures at $OD_{600}$ = 0.5.** (**A**) Nucleic acids bound to TtAgo or TtAgo$^{DM}$. (**B**) Abundance of TtAgo and TtAgo$^{DM}$ and their associated smDNA guides. (**C**) Abundance of smDNA guides bound to TtAgo or TtAgo$^{DM}$, aligned to the chromosome region around the terminus.

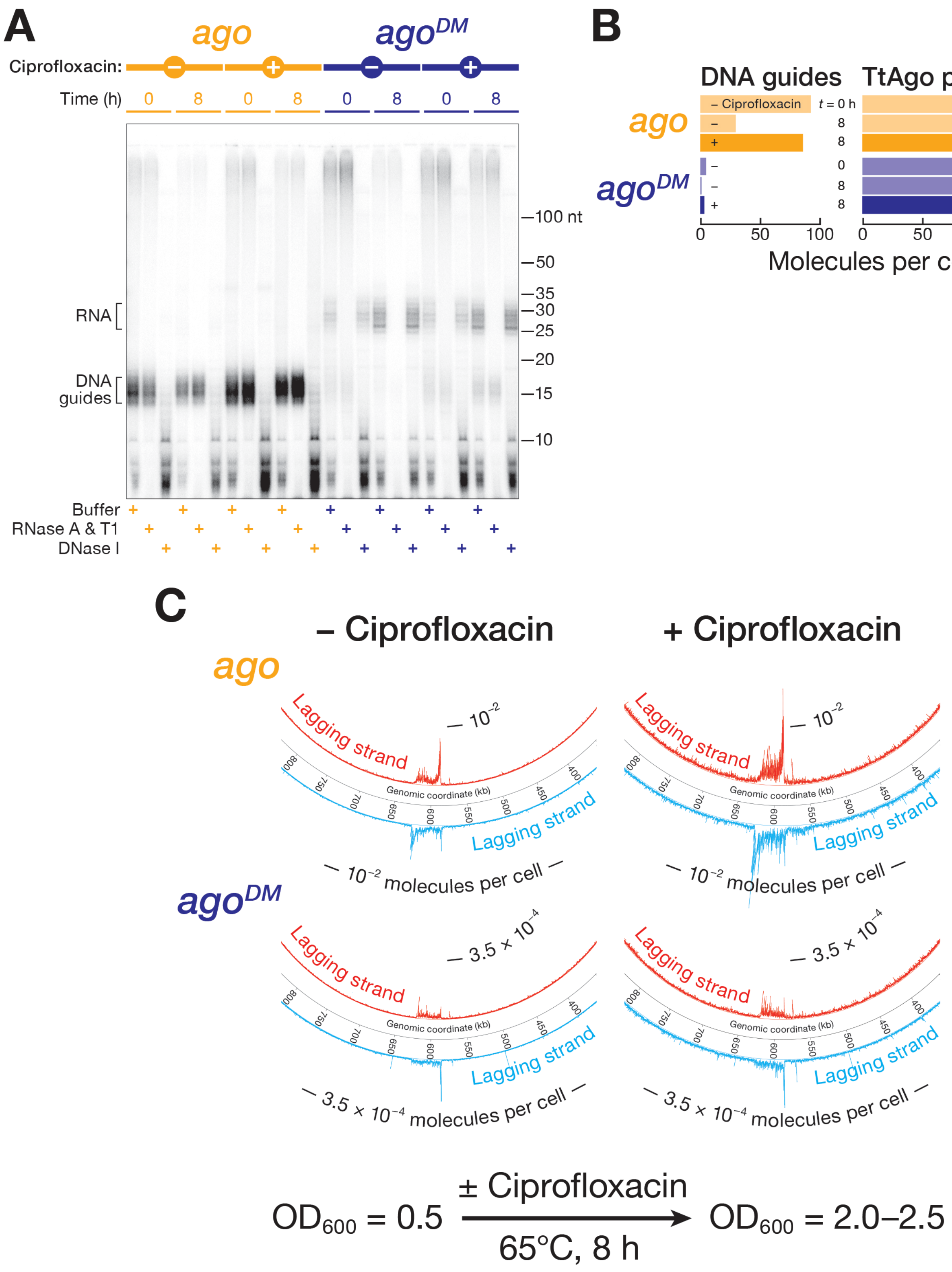
A
ago
agoDM
Ciprofloxacin:
Time (h)
0
8
RNA
DNA guides
100 nt
50
35
30
25
20
15
10
Buffer
RNase A & T1
DNase I
B
DNA guides
TtAgo protein
ago
agoDM
– Ciprofloxacin
t = 0 h
Molecules per cell
C
– Ciprofloxacin
+ Ciprofloxacin
Lagging strand
Genomic coordinate (kb)
10−2 molecules per cell
3.5 × 10−4 molecules per cell
OD600 = 0.5
± Ciprofloxacin
65°C, 8 h
OD600 = 2.0–2.5

**Fig. S5. Proteins bound to TtAgo in kanamycin-marked *T. thermophilus* HB27 strains.** (**A**) Proteins specifically associated with TtAgo at $OD_{600}$ = 0.5 compared to *Δago*. Dashed lines represent FDR = 0.1 (horizontal) and 8× fold enrichment (vertical). Proteins exceeding these cutoffs are highlighted in green (replication or recombination proteins) or black (non-replication or recombination proteins). Proteins not meeting these criteria are grey. (**B**) Silver-stained gel of proteins co-immunoprecipitating with TtAgo or $TtAgo^{DM}$ represented in Fig. 5 and fig. S5A. (**C**) List of all proteins bound to TtAgo with FDR < 0.1 and ≥ 8× fold enrichment (upper left panel of Fig.5).

**A**

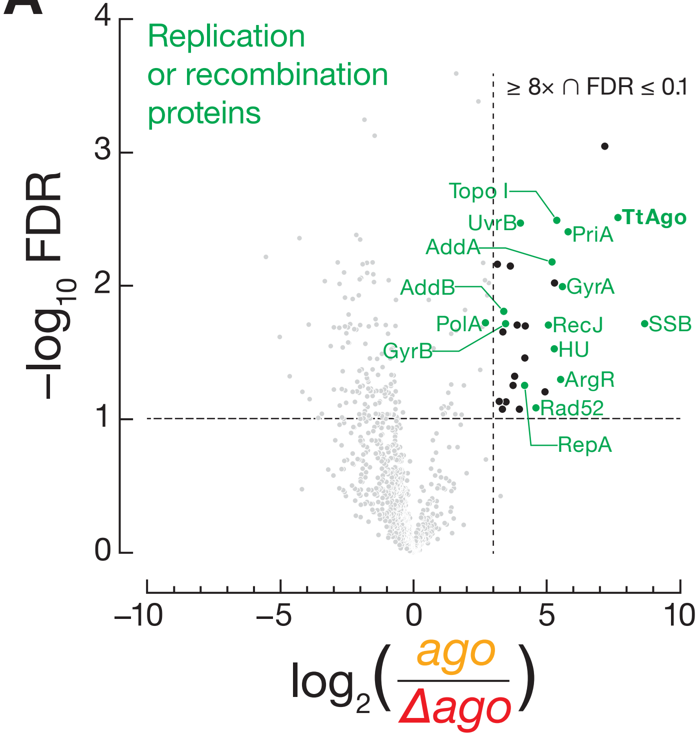


**B**

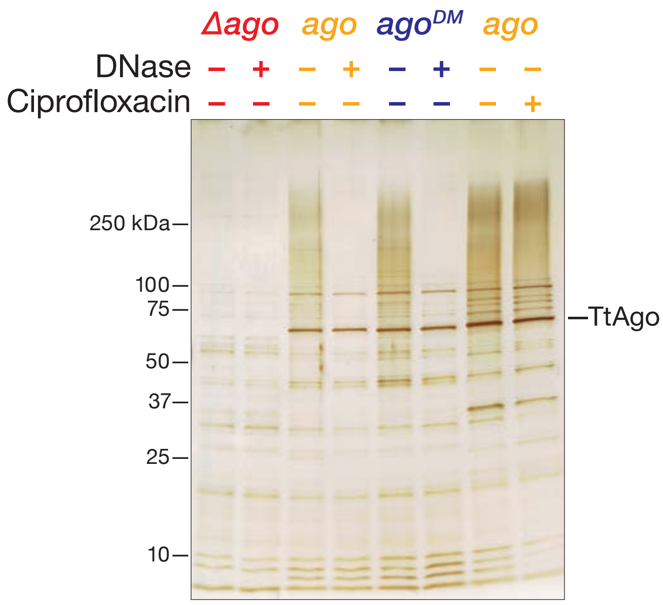


**C**

| Gene | Accession number | Protein | Function | *ago* / *Δago* | FDR |
|---|---|---|---|---|---|
| TT_C1741 | Q72GV7 | ssDNA-binding protein (SSB) | Protect ssDNA | 478 | 0.0228 |
| TT_C0638 | Q72JY0 | AddA | Resect dsDNA breaks, promote recombination | 362 | 0.0048 |
| TT_P0026 | Q746M7 | TtAgo | | 315 | 0.0071 |
| TT_C1942 | Q72GA6 | Primosomal protein N' (PriA) | Restart stalled replication forks | 128 | 0.0042 |
| TT_C0965 | Q72J16 | Glutamine synthetase | Biosynthesis of glutamine | 104 | 0.0012 |
| TT_P0200 | Q745V5 | Uncharacterized | Tetracycline resistance determinant leader peptide | 49 | 0.0288 |
| TT_C0990 | Q72IZ1 | DNA gyrase subunit A (GyrA) | Induce negative supercoiling; decatenation | 49 | 0.0197 |
| TT_C0639 | Q72JX9 | AddB | Resect dsDNA breaks,promote recombination | 45 | 0.0177 |
| TT_C0803 | Q72JG9 | RecJ | ssDNA 5'-to-3' exodeoxyribonuclease | 39 | 0.0290 |
| TT_C0984 | Q72IZ7 | Putative DNA binding protein HU | Protect dsDNA (histone-like) | 32 | 0.0340 |
| TT_C1198 | Q72ID4 | Putative pullulanase | Sugar debranching | 26 | 0.0184 |
| TT_P0198 | Q745V7 | WYL domain-containing protein | Unknown | 24 | 0.0192 |
| TT_C0155 | Q72LA4 | Thiosulfate reductase | Redox reactions | 23 | 0.0687 |
| TT_C1486 | Q72HK1 | Tetratricopeptide repeat-containing protein | Unknown | 21 | 0.0192 |
| TT_C1931 | Q72GB7 | DNA topoisomerase 1 | Relax supercoiled DNA | 20 | 0.0001 |
| TT_C0774 | Q72JJ8 | Polyribonucleotide nucleotidyltransferase (Pnp) | mRNA degradation | 20 | 0.0046 |
| TT_C1194 | Q72ID8 | Arginine repressor (ArgR/XerA) | Arginine biosynthesis; site-specific recombination | 20 | 0.0495 |
| TT_C1923 | Q72GC5 | Rad52/Rad22 superfamily | Repair at dsDNA breaks via recombination | 17 | 0.0763 |
| TT_P0085 | Q746H0 | Replication initiator protein A (RepA) | Initiation of replication | 17 | 0.0549 |
| TT_C0164 | Q72L95 | RNA polymerase sigma factor (SigA) | Transcriptional initiation | 16 | 0.0181 |
| TT_C0162 | Q72L97 | 2'-to-5' RNA ligase superfamily | Putative RNA ligase | 16 | 0.0478 |
| TT_C0124 | Q72LD5 | Glycerate dehydrogenase | Amino acid metabolism | 14 | 0.0297 |
| TT_C0637 | Q72JY1 | Polyphosphate kinase | RNA degradation | 13 | 0.0225 |
| TT_C1531 | Q72HF6 | UvrB | DNA repair | 11 | 0.0241 |
| TT_C0940 | Q72J41 | Putative oxidoreductase | Redox reactions | 11 | 0.0248 |
| TT_C0690 | Q72JS8 | DNA polymerase I | DNA repair; acts at Okazaki fragments | 10 | 0.0210 |
| TT_C0925 | Q72J54 | Ferric uptake regulation protein | Regulation of iron transport proteins | 10 | 0.0986 |
| TT_C1222 | Q72IB0 | DNA gyrase subunit B (GyrB) | Induce negative supercoiling; decatenation | 10 | 0.0196 |
| TT_C1980 | Q72G68 | Glycogen synthase | Synthesis of glycogen | 9 | 0.0294 |
| TT_C1523 | Q72HG4 | ATPase | ATP hydrolysis | 9 | 0.0293 |
| TT_C0121 | Q72LD8 | Phosphoglycerate kinase | Sugar metabolism | 8 | 0.0209 |
| TT_C0928 | Q72J51 | Cell division ATP-binding protein FtsE | Cytokinesis (assembly and stability of septal ring) | 8 | 0.0187 |

**Table S1.** DNA oligonucleotides used in this study.

| Sequence (5′-to-3′) | Description |
|---|---|
| GTG AAT TCG AGC TCG GTA CCC GTC AAA GAA GAA GAT CCC CAC | Δago::htk vector 300 bp upstream TtAgo (F) |
| ATC CGC CGT CAA CGG GTA CCA TCT TGG CTC AGA TTT GCA TAG | Δago::htk vector 300 bp upstream TtAgo (R) |
| ATG CAA ATC TGA GCC AAG ATG GTA CCC GTT GAC GGC GGA T | Δago::htk vector htk gene (F) |
| GGT TCC GCC CCG CTT GGG CAC TGC AGC GTA ACC AAC ATG ATT AAC | Δago::htk vector htk gene (R) |
| TCA TGT TGG TTA CGC TGC AGT GCC CAA GCG GGG CGG AAC C | Δago::htk vector 300 bp downstream TtAgo (F) |
| GGT CGA CTC TAG AGG ATC CCC GCG GGC GAT CTC CTC CTC CG | Δago::htk vector 300 bp downstream TtAgo (R) |
| GAA TTC GAG CTC GGT ACC CGT CAA AGA AGA AGA TCC CCA C | ago$^{DM}$::hyg vector 300 bp upstream TtAgo (F) |
| GTT TTT CCA AGG TGG TTC ATA TCT TGG CTC AGA TTT GCA TAG | ago$^{DM}$::hyg vector 300 bp upstream TtAgo (R) |
| ATG CAA ATC TGA GCC AAG ATA TGA ACC ACC TTG GAA AAA C | ago$^{DM}$::hyg vector TtAgo-$^{DM}$ gene (F) |
| GTG GCG GAT GTC CTG GTG GTC TAA ACG AAG AAG AGC TTT TC | ago$^{DM}$::hyg vector TtAgo-$^{DM}$ gene (R) |
| AAA AGC TCT TCT TCG TTT AGA CCA CCA GGA CAT CCG CCA C | ago$^{DM}$::hyg vector hygromycin (hyg) gene (F) |
| GGT TCC GCC CCG CTT GGG CAA ATA TCT AGA GGA TCC CCT TAT CAC CCG | ago$^{DM}$::hyg vector hygromycin (hyg) gene (R) |
| AAG GGG ATC CTC TAG ATA TTT GCC CAA GCG GGG CGG AAC C | ago$^{DM}$::hyg vector 300 bp downstream TtAgo (F) |
| GGT CGA CTC TAG AGG ATC CCC GCG GGC GAT CTC CTC CTC CG | ago$^{DM}$::hyg vector 300 bp downstream TtAgo (R) |
| CGT TGT AAA ACG ACG GCC AGT GAA TTC GAG CTC GGT ACC CCG CCT GGC GGA CGG GCT T | ago::htk and ago$^{DM}$::htk 300 bp C-terminus TtAgo (F) |
| ATC CGC CGT CAA CGG GTA CCC TAA ACG AAG AAG AGC TTT TCC CGG TCC | ago::htk and ago$^{DM}$::htk 300 bp C-terminus TtAgo (R) |
| AAA AGC TCT TCT TCG TTT AGG GTA CCC GTT GAC GGC GG | ago::htk and ago$^{DM}$::htk vector htk gene (F) |
| GGT TCC GCC CCG CTT GGG CAC TGC AGC GTA ACC AAC ATG ATT AAC | ago::htk and ago$^{DM}$::htk vector htk gene (R) |
| TCA TGT TGG TTA CGC TGC AGT GCC CAA GCG GGG CGG AA | ago::htk and ago$^{DM}$::htk vector 300 bp downstream TtAgo (F) |
| CCA AGC TTG CAT GCC TGC AGG TCG ACT CTA GAG GAT CCC CGC GGG CGA TCT CCT CCT CCG | ago::htk and ago$^{DM}$::htk vector 300 bp downstream TtAgo (R) |
| /5Phos/TGA GGT AGT AGG TTG TAT AGT | Guide for ssDNA and dsDNA and DNA-RNA hybrid binding assay |
| /5Phos/UGA GGU AGU AGG UUG UAU AGU | Guide for ssRNA and dsRNA binding assay |
| /5Phos/CCT CCT CGT CCT CCA A | Guide for stoichiometric binding assays and cleavage assays comparing TtAgo and TtAgo$^{DM}$ |
| /5Phos/GAT CAA CAT TGG AGG ACG AGG AGG ACC T | Target for TtAgo and TtAgo$^{DM}$ cleavage assays |
| /5Phos/CGA GGT AGT AGG TTG TAT AGT | ssDNA guide for cleavage assays |
| CTT TAT ATT TAA ATA ATT TAA TAT ACT ATA CAA CCT ACT ACC TCG TAT AAA TTT TTA AAT AAA TAT TGC ATT CAA GCT TTT AAT TTA ATT AAA TGG CC | AT-rich ssDNA and dsDNA target for cleavage assay |
| CTT TAT ATT TAA ATA ATT TAA TAT ACT ATA C | AT-rich forward primer to make dsDNA target |
| GGC CAT TTA ATT AAA TTA AAA GCT TGA ATG CAA TA | AT-rich reverse primer to make dsDNA target |
| CGC TTA GAC CTA CGC CTG CCA GCA ACT ATA CAA CCT ACT ACC TCG TGC AGA GCC CTT GGG CTG GCC GGC ATT CAA GCT TAC GCA TGG TGG ACC TGG CC | GC-rich ssDNA and dsDNA target for cleavage assay |
| CGC TTA GAC CTA CGC CTG | GC-rich forward primer to make dsDNA target |
| GGC CAG GTC CAC CAT | GC-rich reverse primer to make dsDNA target |

| | |
|---|---|
| /5Phos/TTA CGT GGT CCT GAA T | 16 nt spike-in for smDNA sequencing |
| /5Phos/ATA GGA CCG GCT CTA T | 16 nt spike-in for smDNA sequencing |
| /5Phos/TTA TCT CCA GGG CGC GAA T | 19 nt spike-in for smDNA sequencing |
| /5Phos/TGG AAT TCT CGG GTG CCA AGG/3ddC/ | 3′ adapter for smDNA sequencing |
| /5AmMC6/CCT TGG CAC CCG AGA ATT CCA NNN NNN/3AmMO/ | 3′ bridge for smDNA sequencing |
| GTT CAG AGT TCT ACA GTC CGA CGA TC | 5′ adapter for smDNA sequencing |
| /5AmMC6/NNN NNN GAT CGT CGG ACT GTA GAA CTC TGA AC/3AmMO/ | 5′ bridge for smDNA sequencing |
